# Pick Your Poison: Tetrodotoxin Variants Give Pacific Newts a Potential Leg Up in the Coevolutionary Arms Race with Resistant Garter Snake Predators

**DOI:** 10.64898/2026.05.20.726542

**Authors:** Kait B. Malewicz, Kelly E. Robinson, Anne M. Brown, Christopher S. Jeffrey, Casey S. Philbin, Joel W. McGlothlin, Justin A. Lemkul, Chris R. Feldman

## Abstract

Coevolution proceeds through the evolution of traits that mediate ecological interactions and evolutionary outcomes. In the arms race between toxic Pacific newts (*Taricha*) and their garter snake predators (*Thamnophis*), this interface involves tetrodotoxin (TTX), an antipredator defense that inhibits nerve and muscle function by blocking voltage-gated sodium channels. In response, snakes have evolved TTX-resistant channels, in some cases leading to snake populations that are nearly invulnerable to TTX. For decades, newt TTX has been treated as a single defensive trait, yet TTX occurs as a family of structurally related analogs that may represent alternative defenses against snakes. Here, we characterize TTX analog diversity in all four species of *Taricha* and evaluate how these compounds interact with sodium channels in coevolved garter snakes. Using LC–MS analysis of newt skin secretions, we detected a diverse suite of TTX analogs previously unrecognized in Pacific newts. We then used molecular docking models to evaluate interactions between various TTX analogs and variants of the skeletal muscle channel (Na_V_1.4) that span the range of TTX resistance in garter snakes. We found that some TTX analogs docked better than canonical TTX in resistant snake channels. Notably, we show that 11-deoxy-4-epi-TTX and 11-deoxy-TTX have favorable interactions with hydrophobic amino-acid substitutions in extremely resistant garter snake sodium channels, potentially circumventing predator resistance to canonical TTX. Our results suggest a complex arms race involving multiple newt TTX analogs and multiple snake sodium channel variants. As such, newts may keep pace with snakes by diversifying their arsenal of chemical weapons.

**Significance:** In western North America, some newts have enough toxin in their skin to kill most predators, but coevolution has led to toxin resistance in their garter snake predators that in some cases is so extreme that snakes should be able to withstand more toxin than newts could possibly possess. Here, we explore the possibility that newts may be able to fight back against these resistant predators through the by evolving a diverse repertoire of toxins. We use chemical analyses and computational modeling to show that newts have a wide variety of toxins, some of which are likely to be more effective against resistant snakes, suggesting that coevolution may continue to occur even when snakes become very resistant.

## Introduction

Antagonistic interactions between species, such as those between predator and prey or parasite and host, often drive reciprocal adaptations (Thompson 1994, 2005; Decaestecker et al. 2007; Buckingham and Ashby 2022). In such scenarios, adaptation and counter-adaptation can escalate, leading to arms-race dynamics of ever increasing or extreme coevolutionary traits (Berenbaum and Zangerl 1998; Brodie et al. 2002; Benkman et al. 2003; Reimche et al. 2020). Coevolutionary arms races are mediated by a “phenotypic interface” of interacting traits that determine the outcome of encounters between ecological partners (Brodie and Ridenhour 2003; Thompson 2005; Nuismer et al. 2007). These interface traits are often complex performance phenotypes built from underlying molecular, physiological, and behavioral components (Brodie and Ridenhour 2003). Thus, a complete understanding of coevolution requires not only documenting broad patterns of phenotypic escalation but also identifying the functional traits that mediate the success and failure of ecological interactions (Brodie and Ridenhour 2003; Thompson 2005; Agrawal and Zhang 2021).

The interaction between toxic Pacific newts (*Taricha* spp.) and their garter snake predators (*Thamnophis* spp.) provides a model system for studying antagonistic coevolution (Brodie and Brodie 1999). Pacific newts possess tetrodotoxin (TTX), a poison held in the skin that they can secrete when provoked by predators (Brodie 1968; Stebbins 2003). TTX is a potent neurotoxin that binds to the outer pore of voltage-gated sodium channels (Na_V_), blocking the flow of sodium ions across the cell membrane and arresting action potentials (Noda et al. 1989; Terlau et al. 1991; Hanifin 2010; Abal et al. 2017). Thus, TTX-poisoning from newts disrupts normal neuromuscular function, often resulting in paralysis and death (Brodie 1968). However, three western garter snake species (*T. atratus, T. couchii, T. sirtalis*) have overcome this dangerous defense and can prey on sympatric newts (Brodie and Brodie 1990; Brodie et al. 2002; Brodie et al. 2005; Wiseman and Pool 2007; Greene and Feldman 2009). These three snake species have independently evolved resistance to TTX via specific amino acid substitutions in sodium channels expressed in the periphery, reducing the ability of TTX to bind to these critical proteins and maintaining physiological function in the face of toxin concentrations that would be fatal to most predators (Geffeney et al. 2005; Feldman et al. 2009; Hague et al. 2017; McGlothlin et al. 2016; del Carlo et al. 2024). Work on this system shows extensive geographic variation in both newt TTX levels and snake TTX-resistance, which is mediated primarily via alternative forms of the skeletal muscle channel Na_V_1.4 (Brodie and Brodie 1990; Brodie et al. 2002; Hanifin et al. 2008; Feldman et al. 2010; Stokes et al. 2015; Hague et al. 2017; 2020; Reimche et al. 2020; Gilbert et al. 2023). In general, newt TTX levels and snake TTX resistance are well-matched across populations (Hanifin et al. 2008; Hague et al. 2020; Reimche et al. 2020; Gilbert et al. 2023). However, there are striking mismatches between prey and predator traits in some populations, with some snakes exhibiting such extreme TTX resistance that they cannot be harmed by even the highest levels of TTX in sympatric newts (Brodie et al. 2002; Hanifin et al. 2008; Reimche et al. 2020). At a mechanistic level, these mismatches stem from mutant sodium channels in snakes that are nearly impervious to TTX blockade (Geffeney et al. 2005; Feldman et al. 2009, 2010; Hague et al. 2017; 2020; del Carlo et al. 2024). At an evolutionary level, this pattern suggests that newts may be hampered in their ability to respond to their predators by physiological upper limits to TTX concentrations (Hanifin et al. 2008).

Despite decades of work on this coevolutionary arms race, most research has treated the prey weapon as a simple trait: the amount of canonical TTX in a newt. That simplification has been useful for understanding broad patterns of trait escalation and geographic matching but likely underestimates the chemical complexity of the prey phenotype at the interface of coevolution. TTX is a low molecular weight (∼300 Da) guanidinium toxin that occurs as a family of structurally related compounds (Yasumoto and Yotsu-Yamashita 1996; Yotsu-Yamashita 2001; Hanifin 2010). Multiple salamandrid (newt) species possess canonical TTX as well as a number of similar TTX analogs that differ in stereochemistry, ring structure, and functional groups (Buchwald et al. 1964; Wakely et al. 1966; Mori 1988; Yasumoto et al. 1988; Yotsu et al. 1990; Kotaki and Shimizu 1993; Hanifin et al. 1999; Yotsu-Yamashita and Mebs 2001; Hanifin et al. 2002; Goris and Maeda 2004; Yotsu-Yamashita et al. 2007; Kudo et al. 2012; Yotsu-Yamashita et al. 2012; Kudo et al. 2016; Yotsu-Yamashita et al. 2017; Kudo and Yotsu-Yamashita 2019; Kudo et al. 2020). However, the functional consequences and ecological significance of this analog diversity remain largely unexplored, particularly in *Taricha* species other than *T. granulosa* (Kotaki and Shimizu 1993; Hanifin et al. 1999; Kudo et al. 2020).

Understanding this chemical diversity is important because the phenotypic interface between newts and snakes is ultimately molecular. Tetrodotoxin and analogs act by binding with specific amino acid residues that create and line the outer pore of the sodium channel (Tikhonov and Zhorov 2005; Fozzard and Lipkind 2010; Stevens et al. 2011). The molecular contacts between toxin and channel include polar side chain interactions, backbone-amide interactions, and cation-π interactions that constitute an extensive network of salt bridges and hydrogen bonds that stabilizes and orients TTX in the pore (Tikhonov and Zhorov 2005; Fozzard and Lipkind 2010; Shen et al. 2018). Resistance-conferring substitutions in snake Na_V_ channels include replacements at TTX-binding sites and/or adjacent sites that alter the biophysical environment of the pore (Geffeney et al. 2005; Feldman et al. 2009, 2010, 2012; Hague et al. 2017; del Carlo et al. 2024). However, phenotypic resistance in snakes has only been measured using canonical TTX, as has resistance in the tissues (skeletal muscle) and sodium channels themselves (Geffeney et al. 2002, 2005; McGlothlin et al. 2016; Reimche et al. 2022; del Carlo et al. 2024). Thus, the effects of other naturally occurring TTX analogs remain largely unknown. It is possible that the evolutionary response of newts involves more than just changes in the quantity of TTX, extending to diversification of the structure of TTX itself. If some naturally occurring TTX analogs interact more favorably with resistant snake channels than canonical TTX, then chemical diversification may represent an underappreciated aspect of the phenotypic interface of coevolution.

The use of chemically modified forms of TTX that are more effective against TTX-resistant channels could offer a pathway for renewed escalation of the coevolutionary arms race between newts and snakes. Here, we test this hypothesis by characterizing TTX analog diversity in *Taricha* and evaluate how alternative TTX structures are predicted to interact with the Na_V_1.4 channels of the three *Thamnophis* species, spanning the full range of resistance phenotypes. Specifically, we (1) characterize the diversity and relative abundance of TTX analogs in all four Pacific newt species (table 1) using liquid chromatography–mass spectrometry (LC–MS) to determine whether *Taricha* possess a broader collection of TTX compounds than has been appreciated, (2) evaluate whether some TTX analogs might bind to snake sodium channels more favorably than canonical TTX, especially in extremely TTX-resistant Na_V_ (fig. 1), and (3) characterize the molecular interactions that might be responsible for more the favorable contacts between newt TTX analogs and snake sodium channels. These complementary approaches allow us to explore whether selection has pushed newts to exploit a diversity of TTX analogs to overcome the resistance mechanisms of their snake adversaries, refining our understanding of the coevolutionary interface in this classic arms-race system.

**Table 1.** Summary information on newts sampled for neurotoxins.

| Newt Species<sup>a</sup> | Newt Locality | Sample Size (n) | Sympatric Snake Predator<sup>b</sup> |  |  | Latitude, Longitude |
| --- | --- | --- | --- | --- | --- | --- |
|  |  |  | *T. sirtalis* | *T. atratus* | *T. couchii*<sup>c</sup> |  |
| *T. granulosa*<br>(n = 10) | North Mill Creek, Mendocino Co., CA | 2 | ✓ | ✓ |  | 38.9062, -123.3256 |
|  | Galbreath Preserve, Mendocino Co., CA | 4 | ✓ | ✓ |  | 38.8508, -123.2660 |
|  | Mitsui Ranch Preserve, Sonoma Co., CA | 4 | ✓ | ✓ |  | 38.3246, -122.5799 |
| *T. rivularis*<br>(n = 10) | Galbreath Preserve, Mendocino Co., CA | 5 | ✓ | ✓ |  | 38.8682, -123.2526 |
|  | Little Sulphur Creek, Sonoma Co., CA | 5 |  | ✓ |  | 38.7642, -122.8166 |
| *T. sierrae*<br>(n = 10) | Fiddle Creek, Sierra Co., CA | 5 | ✓ |  | ✓ | 39.5216, -120.9931 |
|  | Canyon Creek, Nevada Co., CA | 5 |  |  | ✓ | 39.4429, -120.6581 |
| *T. torosa*<br>(n = 10) | Mitsui Ranch Preserve, Sonoma Co., CA | 3 | ✓ | ✓ |  | 38.3246, -122.5799 |
|  | Bear Creek, Contra Costa Co., CA | 3 | ✓ | ✓ |  | 36.2011, -118.7609 |
|  | Big Falls Canyon, San Luis Obispo Co., CA | 4 | ✓ | ✓ |  | 35.2623, -120.5118 |

<sup>a</sup>*Taricha*.
<sup>b</sup>*Thamnophis*.
<sup>c</sup>*T. couchii* also interacts with *T. granulosa* and *T. torosa* at the northern and southern ends of its distribution, respectively, but we did not sample newts from those areas for this study.

**Fig. 1.**
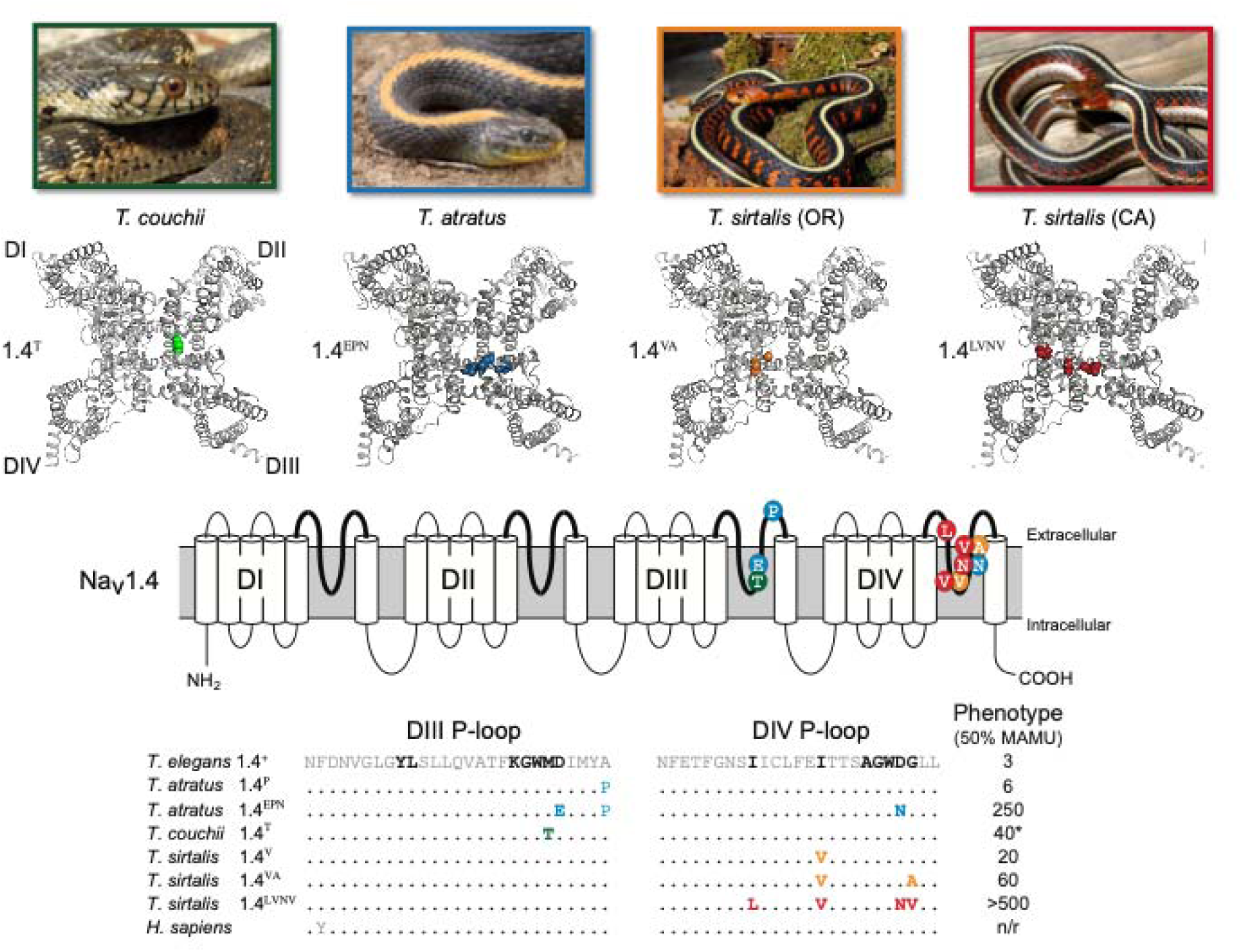
Three species of garter snakes (Thamnophis) have evolved resistance to tetrodotoxin (TTX), the defensive poison of Pacific newts (Taricha). Tetrodotoxin binds to the outer pore of voltage-gated sodium channels (center of 3D ribbon diagram), blocking the movement of Na^+^ through the channel. Some populations of T. couchii, T.atratus, and T. sirtalis possess skeletal muscle sodium channels (Na_v_1.4) with replacements in the outer pore (P-loops) of Domains III and IV, conferring differing levels of TTX resistance to the channel, and thus to snakes. Amino acid replacements are color coded by species (or population) and shown in 3D and 2D representations of Na_v_1.4, and on amino acid alignment. Reference DIII and DIV P-loop sequence of T. elegans at top of alignment exhibits the ancestral TTX-sensitive channel (Na 1.4^+^) seen across nearly all garter snake species. Whole animal resistance is the dose of TTX required to reduce sprint speed of a snake by 50%, shown in Mass Adjusted Mouse Units (50% MAMU). Photos (left to right): G. Nafis, R Sikola, S. Bol, G Nafis.

## Results

### TTX Analogs in Newts

We identified nine compounds as TTX and TTX analogs (table 2). The analogs we detected include epi-TTX, anhydro-TTX, deoxy-TTX and other TTX species. However, our mass spectrometric methods were unable to parse known epimers and deoxy analogs. Previous work had only identified 6-epi-TTX and 11-deoxy-TTX in *T. granulosa* (Kotaki and Shimizu 1993; Hanifin et al. 1999; Kudo et al. 2020). We detected these analog types and others in all four species of *Taricha*. In addition, we detected a TTX analog we putatively identify as 11-oxotetrodotoxin (compound **5**) that has not previously recorded in *Taricha* (table 2). This TTX analog was produced synthetically from tetrodotoxin (Wu et al. 1996) before being discovered naturally in the eastern newt, *Notophthalmus viridescens* (Yotsu-Yamashita and Mebs 2003). We assumed that ionization efficiency of TTX analogs was similar to TTX, allowing for comparisons of relative analog concentrations as previously reported in pufferfish (Park et al. 2024). Using ANOVA, we found that newt species differed in the relative abundance of the nine TTX analogs (F_30,396_ = 5.50; p < 0.001). Our *post hoc* analyses revealed that newt species primarily differed in three compounds (numbering follows table 2; fig. 2, 3): anhydro-TTX (**10** or **11**); deoxy-TTX (**6** or **7**); and canonical TTX (**1**) (fig. 3). For all but one species, canonical TTX was the most abundant alkaloid we detected, and our sample of *T. rivularis* contained the highest relative amounts of TTX, followed by *T. granulosa*. However, our sample of *T. sierrae* possessed higher amounts of deoxy-TTX (**6** or **7**), while our sample of *T. torosa* possessed low relative abundances of all TTX analogs. We found that species also differed in their levels of analog diversity (p < 0.001, F_3,36_ = 13.28). Our *post hoc* analysis of Simpson’s diversity revealed that *T. granulosa* (D = 0.59) and *T. rivularis* (D = 0.52) had similar and higher mean TTX analog diversity compared to *T. sierrae* (D = 0.24) and *T. torosa* (D = 0.34) with similar and lower levels of analog diversity.

**Table 2.** Detected compounds in newt skin secretions, along with known TTX analogs from the literature.

| Exp. Mass (Da) | Theo. Mass (Da) | Diff (ppm) | RT (min) | Chemical Formula | TTX Analog Type | Newt Species<sup>a</sup> |  |  |  | Cpd. No. | Literature TTX Analogs | Literature Newt Species<sup>b</sup> |
| --- | --- | --- | --- | --- | --- | --- | --- | --- | --- | --- | --- | --- |
| 319.1023 | 319.1016 | -2.2 | 1.48 | C<sub>11</sub>H<sub>17</sub>N<sub>3</sub>O<sub>8</sub> | TTX | Tg | Tr | Ts | Tt | 1 | TTX | *Taricha*,<br>*Notophthalmus*,<br>*Cynops*, *Triturus*<br>(1-12) |
| 319.1021 | 319.1016 | -1.6 | 1.11 | C<sub>11</sub>H<sub>17</sub>N<sub>3</sub>O<sub>8</sub> | epi-TTX | Tg | Tr |  |  | 2 | 4-epi-TTX□ | *Cynops ensicauda*<br>(1);<br>*T. granulosa* (3) |
|  |  |  |  |  |  |  |  |  |  | 3 | 6-epi-TTX□ | *Taricha*,<br>*Notophthalmus*,<br>*Cynops*, *Triturus*<br>(1-12) |
|  |  |  |  |  |  |  |  |  |  | 4 | 8-epi-TTX | *Cynops ensicauda*<br>(7) |
| 335.0967 | 335.0965 | -0.6 | 1.77 | C<sub>11</sub>H<sub>17</sub>N<sub>3</sub>O<sub>9</sub> | oxo-TTX | Tg | Tr | Ts | Tt | 5 | 11-oxo-TTX | *Notophthalmus* (6) |
| 303.1055 | 303.1066 | 3.6 | 1.72 | C<sub>11</sub>H<sub>17</sub>N<sub>3</sub>O<sub>7</sub> | deoxy-TTX | Tg | Tr | Ts | Tt | 6 | 11-deoxy-TTX | *Cynops ensicauda*<br>(1);<br>*T. granulosa* (3, 5) |
|  |  |  |  |  |  |  |  |  |  | 7 | 11-deoxy-4-epi-TTX | *Cynops ensicauda*<br>(1) |
| 303.1070 | 303.1066 | -1.3 | 1.13 | C<sub>11</sub>H<sub>17</sub>N<sub>3</sub>O<sub>7</sub> | deoxy-TTX | Tg | Tr |  |  | 8 | 1-hydroxy-8-epi-5,11-dideoxy-TTX | *T. granulosa* (2, 3) |
|  |  |  |  |  |  |  |  |  |  | 9 | 11-deoxy-TTX (lactone form) | *T. granulosa* (3) |
| 287.1116 | 287.1117 | 0.3 | 1.47 | C<sub>11</sub>H<sub>17</sub>N<sub>3</sub>O<sub>6</sub> | dideoxy-TTX | Tg |  |  | Tt | 10 | 8-epi-5,11-dideoxy-TTX | *T. granulosa* (2) |
| 287.1118 | 287.1117 | -0.3 | 2.13 | C<sub>11</sub>H<sub>17</sub>N<sub>3</sub>O<sub>6</sub> | dideoxy-TTX | Tg |  | Ts |  | 11 | 5,11-dideoxy-TTX□ | *T. granulosa* (2) |
| 301.0910 | 301.0910 | 0.0 | 1.58 | C<sub>11</sub>H<sub>15</sub>N<sub>3</sub>O<sub>7</sub> | anhydro-TTX | Tg | Tr | Ts | Tt | 12 | 4,9-anhydro-TTX | *Cynops ensicauda*;<br>*T. granulosa* (3,4) |
|  |  |  |  |  |  |  |  |  |  | 13 | 4,9-anhydro-6-epi-TTX | *Cynops ensicauda*<br>(1) |
| 285.0960 | 285.0961 | 0.4 | 6.87 | C<sub>11</sub>H<sub>15</sub>N<sub>3</sub>O<sub>6</sub> | anhydro, deoxy-TTX | Tg | Tr | Ts |  | 14 | 4,9-anhydro-11-deoxy-TTX□ | *Cynops ensicauda*<br>(1) |

<sup>a</sup>Tg: *Taricha granulosa*, Tr: *T. rivularis*, Ts: *T. sierrae*, Tt: *T. torosa*
<sup>b</sup>References: 1. Yasumoto et al. (1988); 2. Kudo et al. (2020); 3. Kotaki and Shimizu (1993); 4. Hanifin et al. 2002; 5. Yotsu et al. (1990); 6. Yotsu-Yamashita and Mebs (2003); 7. Kudo and Yotsu-Yamashita (2019); 8. Kudo et al. (2016); 9. Hanifin et al. (1999); 10. Yotsu-Yamashita et al. (2007); 11. Yotsu-Yamashita et al. (2012); 12. Yotsu-Yamashita et al. (2017).

**Fig. 2.**
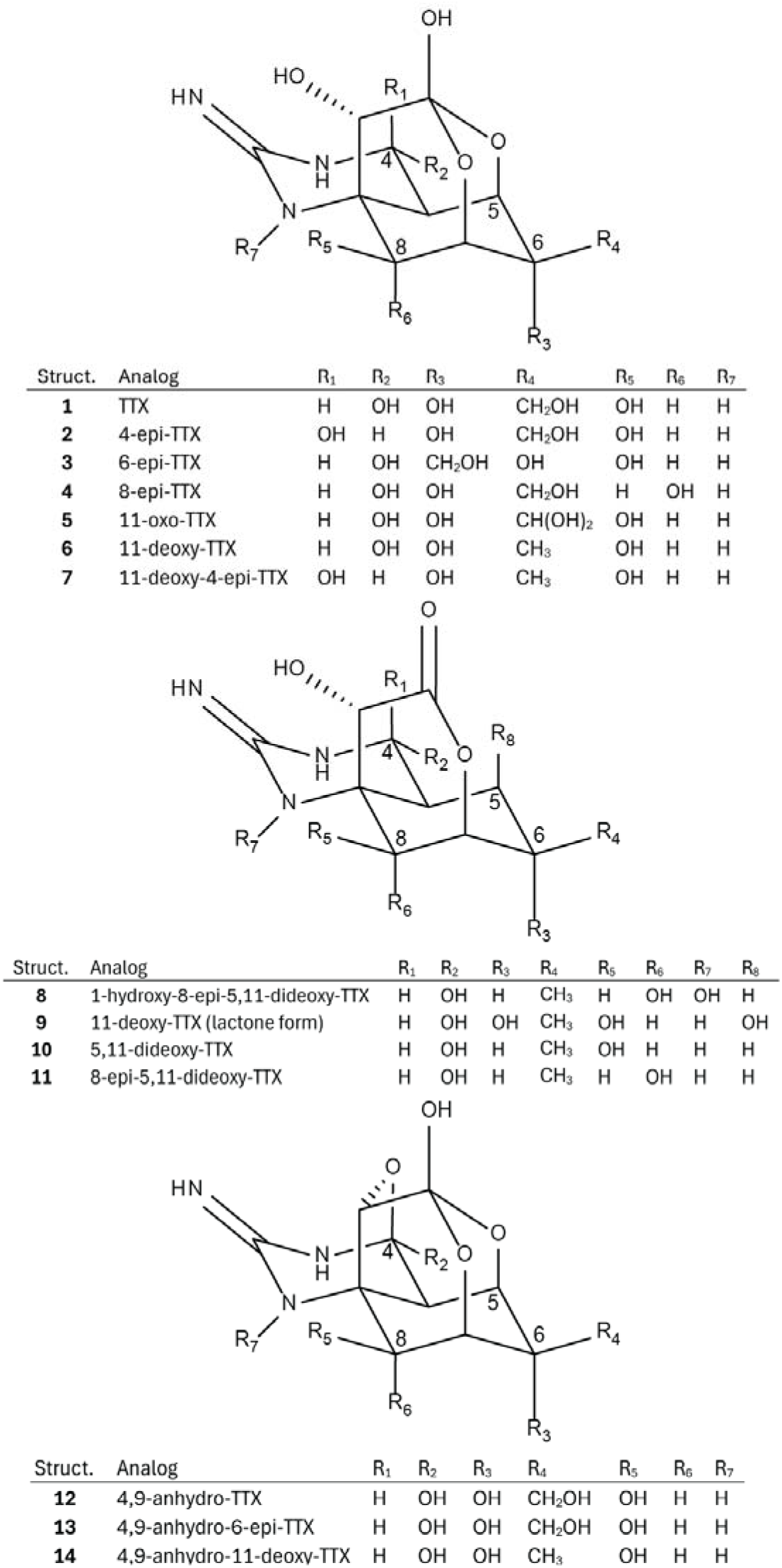
Tetrodotoxin (TTX) analogs reported from newts. Structures of TTX analogs previously detected in newts (of the family Salamandridae) in the literature (see table 2). Compounds numbered for quick reference. Tables below each structure indicate the substitute groups at variable positions in the TTX scaffold. Note that compounds 1-4, 6-9, 12 and 13 were used for molecular docking, and in the oxyanionic form with the C10-oxygen deprotonated.

**Fig. 3.**
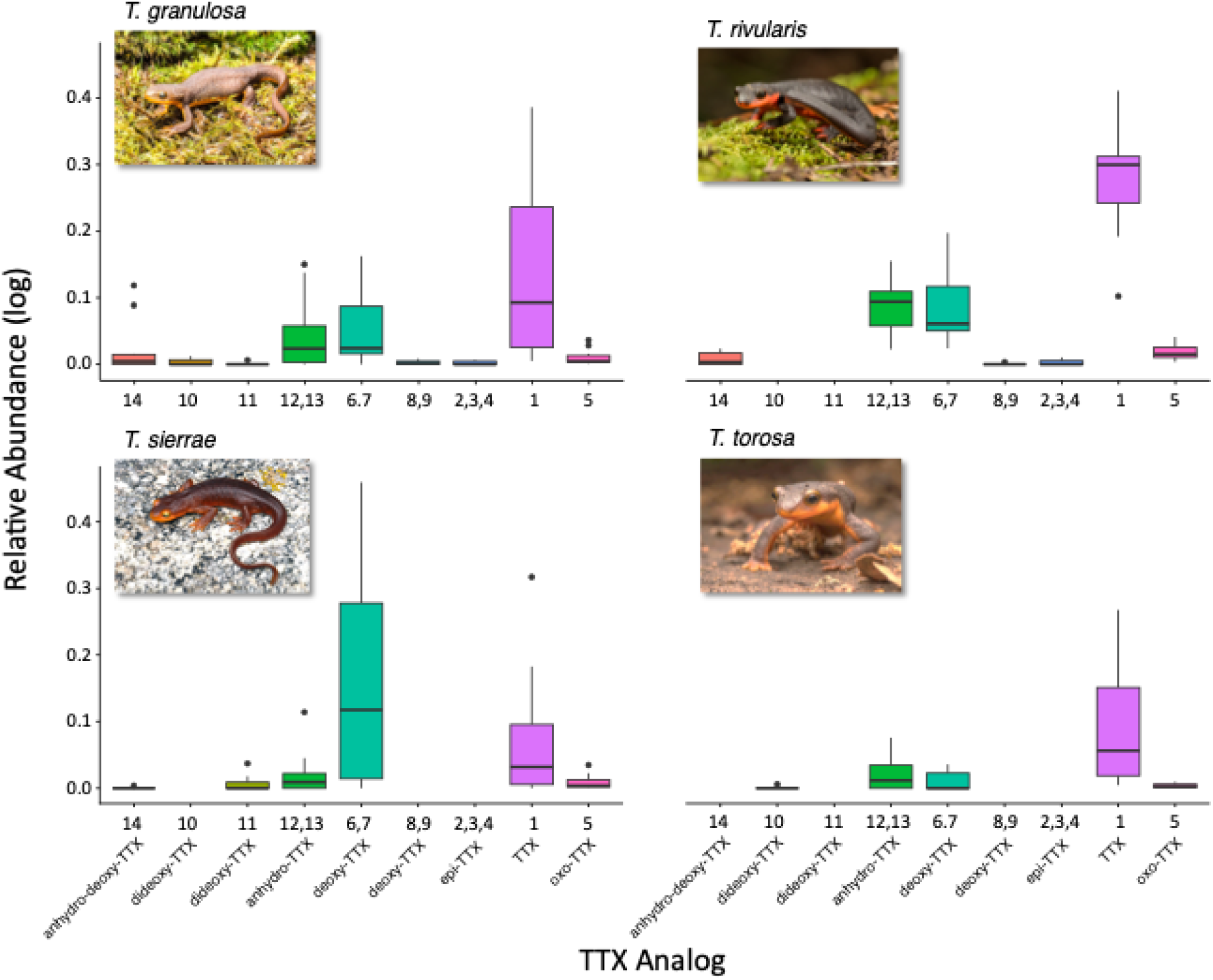
Relative abundance of TTX analogs Pacific newt (Taricha) species. Boxplots depict the log-transformed (log_10_[x] +1) relative abundance of TTX analogs detected in each newt species. Analogs organized by mass (lowest to highest). Note that our mass spectrometric methods could not fully resolve all known epimers and deoxy analogs, thus TTX analogs are reported as TTX analog type and compound number (see table 2; Fig. 1). Photos (top): J Clare, J Fuller; (bottom): RW Hansen, A Sorokin.

The guanidinium group engaged in charge-charge interactions with carboxylate groups of acidic residues in DI and DII, including D375 and E704. E378 formed a hydrogen bond with the C4 hydroxyl group of TTX, while Y376 contributed a cation-π interaction with the guanidinium group. Additional electrostatic contacts were observed with E704, which interacted with both the guanidinium nitrogens and the C8 hydroxyl group, and with E701, which coordinated the C9 OH group via both side chain oxygens, consistent with prior structural observations.

Several backbone-mediated interactions that interact with the TTX oxyanion were also recovered. The backbone carbonyl oxygen of F1060 formed a hydrogen bond with the C9 hydroxyl group, while the backbone amide groups of G1062, W1063, and L1064 served as hydrogen bond donors with both the C10 oxyanion and the ether oxygen between C7 and C10. Notably, the W1063 indole N atom at this position also served as a hydrogen bond acceptor for the C9 hydroxyl group. The docked pose also predicted an additional backbone interaction with K1061 that contacts the oxyanion. Further stabilization was provided by the backbone amide interactions between G1354 and the C4 hydroxyl group as well as hydrogen bonding between D1356 and the C11 hydroxyl group. Together, this elaborate network of side chain and backbone interactions collectively reproduces the experimentally resolved TTX binding mode. As such, our docking approach was validated and serves as a robust baseline for TTX-Na_V_ interactions. Moreover, this interaction framework provides a reference against which deviation in binding geometry and contact composition can be assessed in subsequent docking analyses of other sodium channels and TTX analogs, enabling direct comparison of conserved versus divergent determinants of TTX sensitivity.

### Docking TTX and TTX Analogs to the Cockroach Na_V_PaS

To validate our docking protocol, we first redocked TTX to Na_V_PaS (PDB: 6A95), the cockroach channel that was experimentally resolved via cryo-EM with TTX bound to the pore (Shen et al. 2018). The top-ranked pose recovered the experimentally determined TTX binding mode, including both the overall ligand position within the pore and the specific residue interactions observed in the cryo-EM structure (fig. 4a). This strong agreement indicates that our docking protocol reliably captures the native TTX-Na_V_PaS channel interactions and is suitable for comparative analyses across *Thamnophis* Na_V_1.4 variants and newt TTX analogs.

**Fig. 4.**
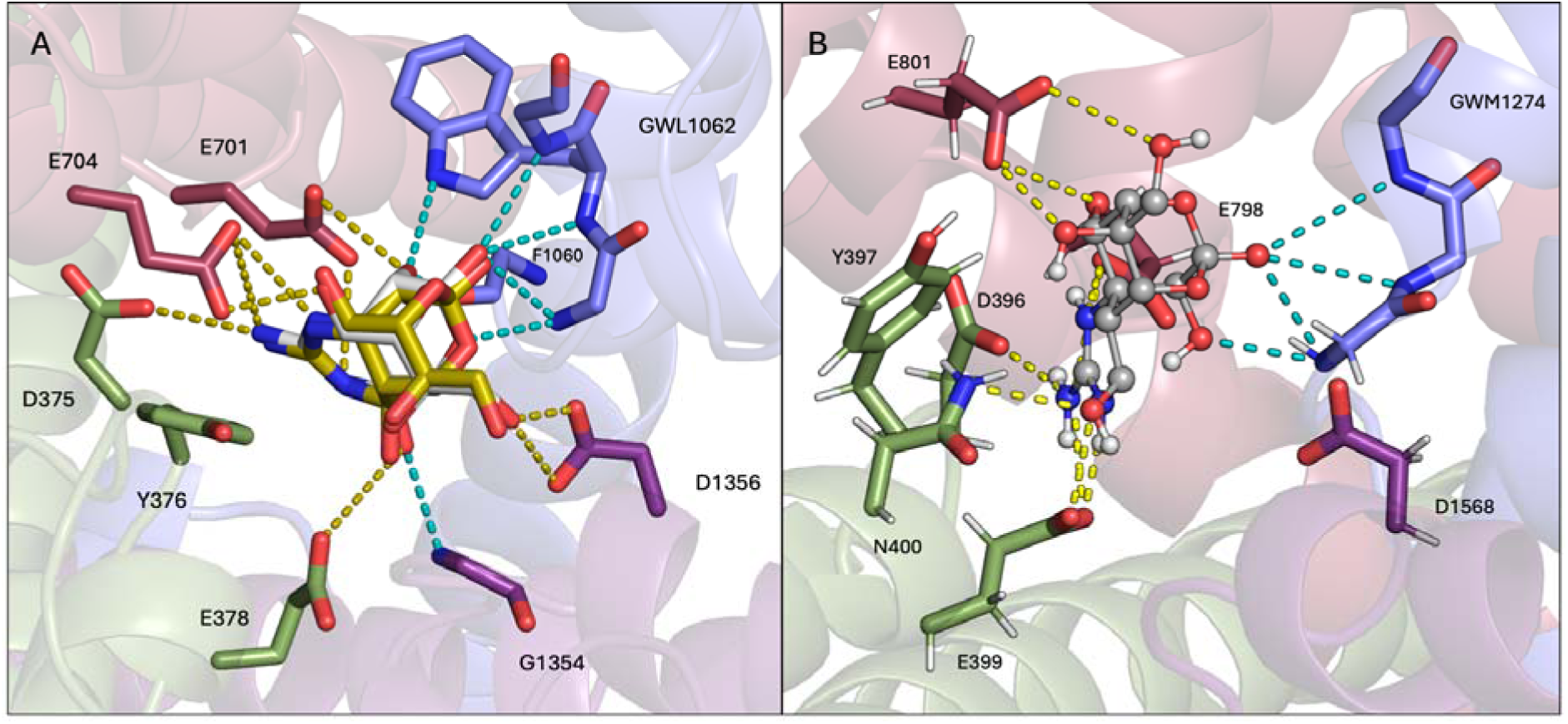
a) Redocked pose of TTX. The first redocked pose of TTX (yellow carbon atoms) is superimposed onto the binding pose from the cryo-EM structure (light gray carbon atoms). Specific interactions are indicated by dashed lines, with backbone interactions in blue and polar side chain interactions in yellow. The first redocked TTX pose reproduced all major contacts previously reported for the sodium channel of the cockroach (Na_V_PaS). The guanidinium group engaged in charge-charge interactions with carboxylate groups of acidic residues in DI and DII, including D375 and E704. E378 formed a hydrogen bond with the C4 hydroxyl group of TTX, while Y376 contributed a cation-π interaction with the guanidinium group. Additional electrostatic contacts were observed with E704, which interacted with both the guanidinium nitrogens and the C8 hydroxyl group, and with E701, which coordinated the C9 OH group via both side chain oxygens, consistent with prior structural observations (Shen et al. 2018). b) First pose of TTX (gray carbon atoms) in Na_V_1.4. Specific interactions are indicated by dashed lines, with backbone interactions in blue and polar side chain interactions in yellow.

### Canonical TTX Contacts in Na_V_1.4 Pore

The first pose of TTX in Na_V_1.4 recovered most of the canonical interactions seen in the cockroach channel (fig. 4b), even though it is slightly rotated relative to Na_V_PaS (fig. S3). The C6 hydroxymethyl group is rotatable and if we performed dynamics, we could observe a structure that better aligns the first pose of Na_V_1.4 to Na_V_PaS. DI contributes to the positioning of the guanidium group via cation-π interactions from Y397 and hydrogen bond donors in D396 and E399. The C4 hydroxyl shares a hydrogen bond partner with E399 as it does in Na_V_PaS but serves as an acceptor for a hydrogen bond with N400 that is a non-interacting aspartate in Na_V_PaS. DII contributes an additional salt bridge to the guanidium group via E798, compared to Na_V_PaS. E801 may engage in hydrogen bonding with any of the C6, C8, and C11 hydroxyl groups. This positioning slightly deviates from that in Na_V_PaS, which has both DII glutamate residues interacting with the guanidinium and either the C9 (Na_V_PaS: E701) or C8 (Na_V_PaS: E704) hydroxyl groups. DIII contains the backbone amides from the same three residues interacting with the oxyanion, G1274, W1275, and M1276. The W1275 amide N does not interact with the oxyanion as in Na_V_PaS, but the G1274 amide engages in a bifurcated hydrogen bond with the C9 hydroxyl group. Unlike D1356 in Na_V_PaS, the D1568 side chain does not interact with TTX in the top pose. The DIV G1354 also does not interact with TTX as the analogous residue in Na_V_PaS does.

### TTX Analog Docking Scores

Our approach sought to distinguish how different functional group properties of TTX analogs interact with snake Na_V_1.4 variants compared to the canonical structure of TTX. On average, the 11-deoxy analogs of TTX interacted more favorably than did canonical TTX across most of the snake sodium channel variants (fig. 5). The 4-epi and 4,9-anhydro analogs were generally comparable to the canonical TTX docking score but docking improved when 4-epi combined with 11-deoxy modification. The 8-epi and 6-epi analogs were weaker predicted binders than TTX, even when 6-epi was combined with the 4,9-anhydro modification. The lactone versions of TTX (5,11-dideoxy and 11-deoxy lactone) had the weakest docking scores on average across all TTX species, including 11-deoxy in lactone form.

**Fig. 5.**
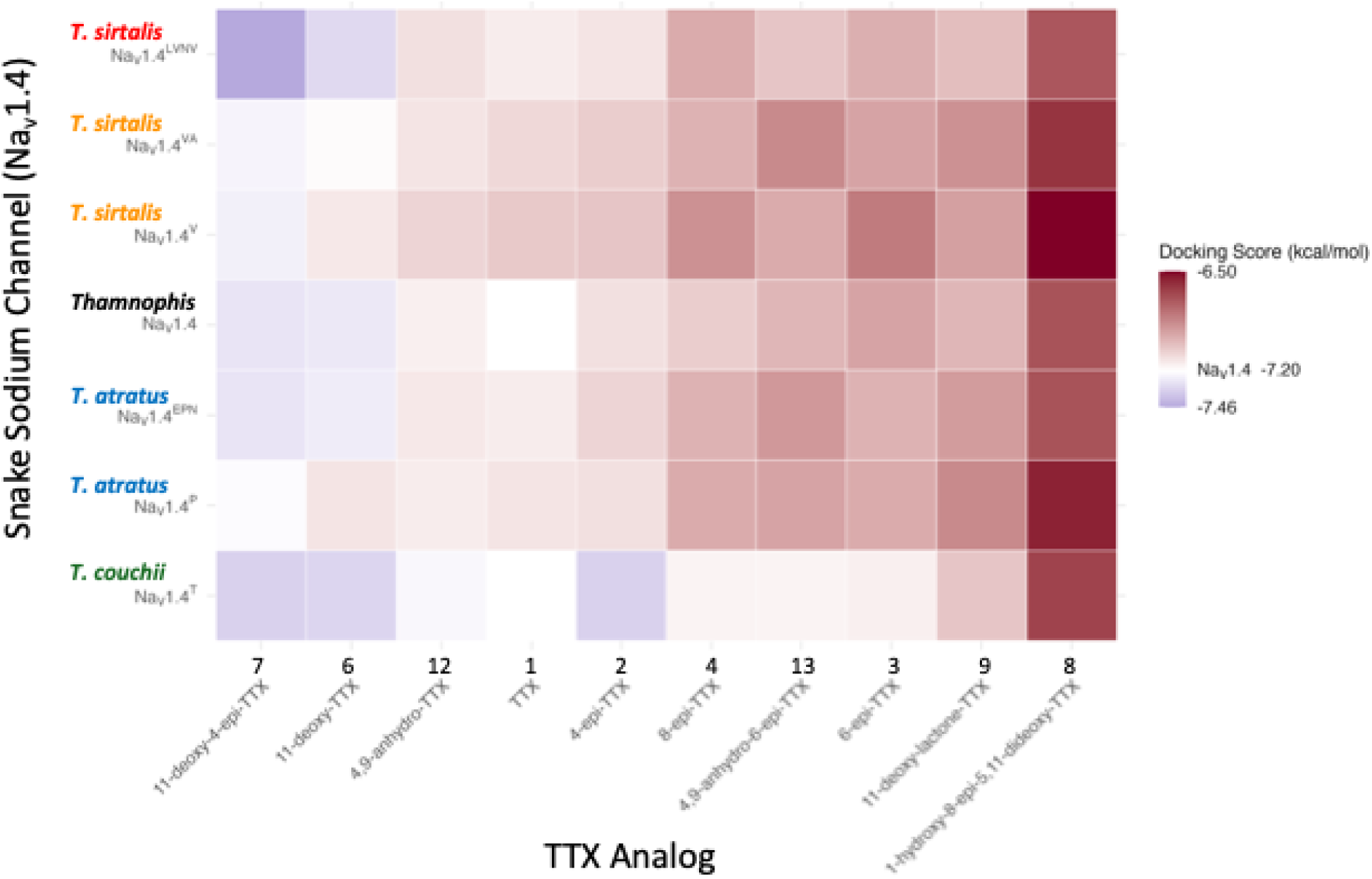
Docking scores of TTX analogs in the sodium channel variants (Na_V_1.4) found in the three snake species engaged with newts. Docking scores (kcal/mol) are averages for each of the nine poses generated in each docking run. Heatmap arranged from highest TTX analog docking in the sodium channel pore (cooler blue colors) to lowest docking (warmer red colors) and scaled to mean docking score (white; −7.20 kcal/mol) of canonical TTX in the ancestral Thamnophis sodium channel (Na 1.4^+^) as a point of reference.

### TTX Contacts in the Extremely Resistant Na_v_1.4^LVNV^ and Sensitive Na_v_1.4^+^ Channels

To further investigate the nature of the docked poses, we generated interaction fingerprints to characterize different classes of interactions for each TTX analog in the extremely resistant *T. sirtalis* channel (Na_V_1.4^LVNV^) and the sensitive snake channel (Na_V_1.4^+^). In both channel variants, Y397, E399, N400, E798, E801, K1273, and G1274 appeared to be important for binding for all analogs (fig. S4) and represent the conserved interaction scaffold for TTX. Despite this shared framework, interaction fingerprints revealed systematic differences between channels. Several residues, including M1276, G1566, and N/D1568, had frequent contacts in both channels but were more pronounced in the Na_V_1.4^LVNV^ channel (fig. S4). Overall, the TTX analogs formed fewer contacts with D396, G1556, and V/G1569 in Na_V_1.4^+^ compared to Na_V_1.4^LVNV^

### Interactions between the best scoring TTX analog in Na_V_1.4^+^ and Na_V_1.4^LVNV^

11-deoxy-4-epi-TTX had the best average score across most of the Na_V_ channels. Notably, the interaction between this variant and the channel with highest resistance to canonical TTX, Na_V_1.4^LVNV^, exhibited the highest average docking score of any channel-analog pair we examined (fig. 5). To investigate what interactions may be contributing to this outcome, we compared interactions at sites 1568 and 1569 in Na_V_1.4^+^ and Na_V_1.4^LVNV^. These are the two of the four sites that are different across the pore of these channels that also have the potential to interact with TTX analogs. Out of all nine poses of 11-deoxy-4-epi-TTX in Na_V_1.4^+^ (fig. S7), five of them involved the guanidium group making at least one contact with a negatively charged DI or DII residue, making these poses suitable for further investigation of intermolecular interactions (fig. 6). In Na_V_1.4^LVNV^, seven out of the nine poses (fig. S5) meet the same criteria of guanidium-DI/DII contacts and were investigated further (fig. 7). Site 1568 is an aspartate in Na_V_1.4^+^ and an asparagine in Na_V_1.4^LVNV^, so both channels have the opportunity for hydrogen bonding and induced dipole interactions, but the negative charge on the aspartate lacks the ability to interact with oxyanion that Na_V_1.4^LVNV^ exhibits in pose 3 (fig. 7c). Otherwise, both channels have site 1568 interacting with the C4 hydroxyl in pose 1 (fig. 6a, 7a), the C6 hydroxyl (fig. 6c, 6d, 7d, 7g), and the C10-C5 ether oxygen (fig. 6b, 6e, 7e). These favorable hydrogen bonds position the C11 methyl group to interact with position 1569; however, since this site is a glycine in Na_V_1.4^+^, there are no interactions available to the C11 methyl group (fig. 6). In Na_V_1.4^LVNV^, site 1569 is a valine, so when the C-11 methyl group is positioned towards it, there are favorable hydrophobic interactions that are not available to the Na_V_1.4^+^ 1569 glycine. Four of the seven poses considered for Na_V_1.4^LVNV^ have direct hydrophobic interactions with V1569 (fig. 7b, d-f), and an additional pose makes hydrophobic contact with C7 and C8 CHs (fig 7g). While an aspartate or an asparagine at site 1568 offers similar contacts with 11-deoxy-4-epi-TTX, a valine at site 1569 better accommodates interactions with the hydrophobic C-11 methyl group than a non-interacting glycine.

**Fig 6.**
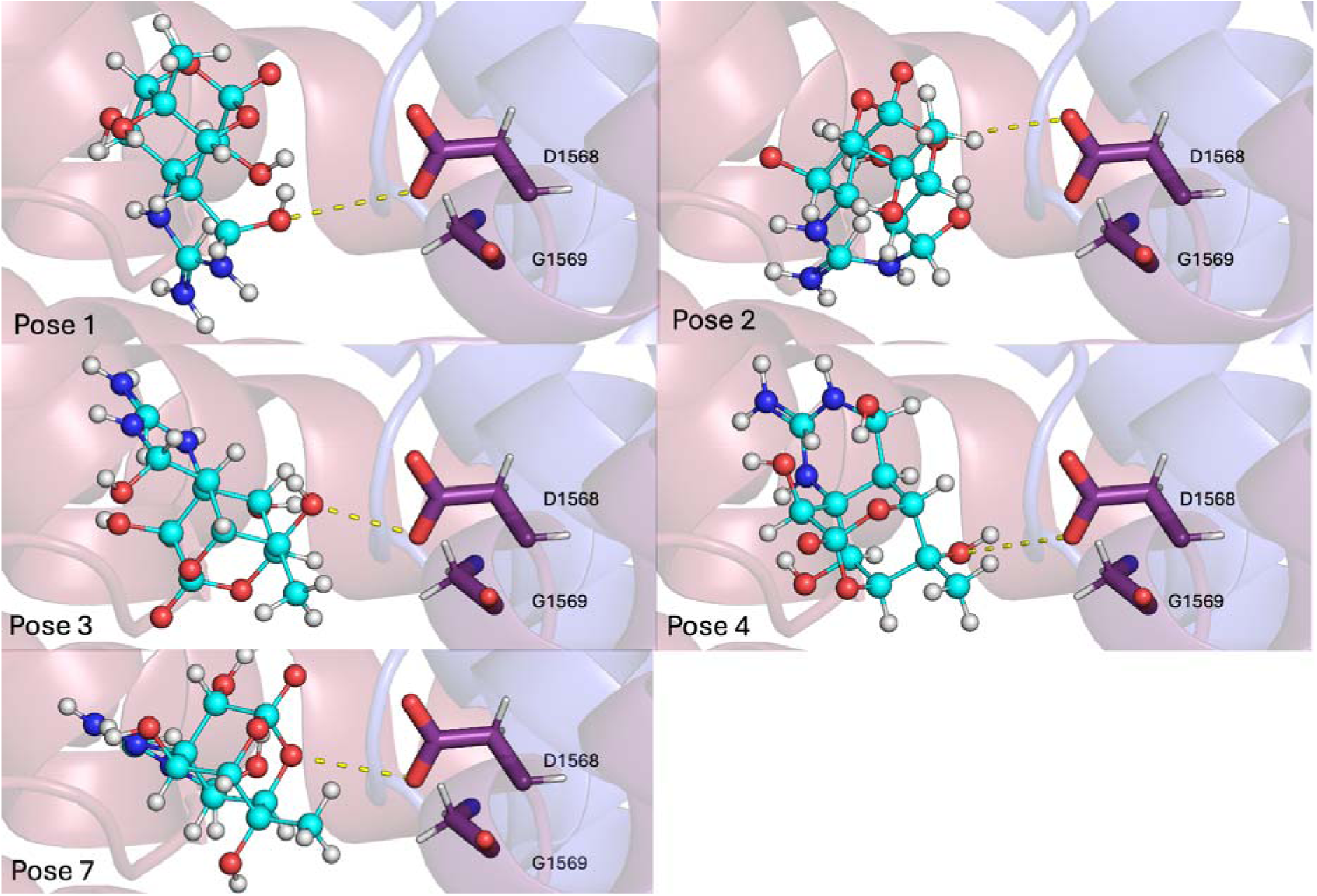
Selected poses (1, 2, 3, 4, 7) of the highest scoring analog, 11-deoxy-4-epi-TTX (cyan, ball and stick) in Na_V_1.4. D1568 and G1569 in Domain IV are shown as sticks with TTX-D1568 interactions shown as dashed lines.

**Fig 7.**
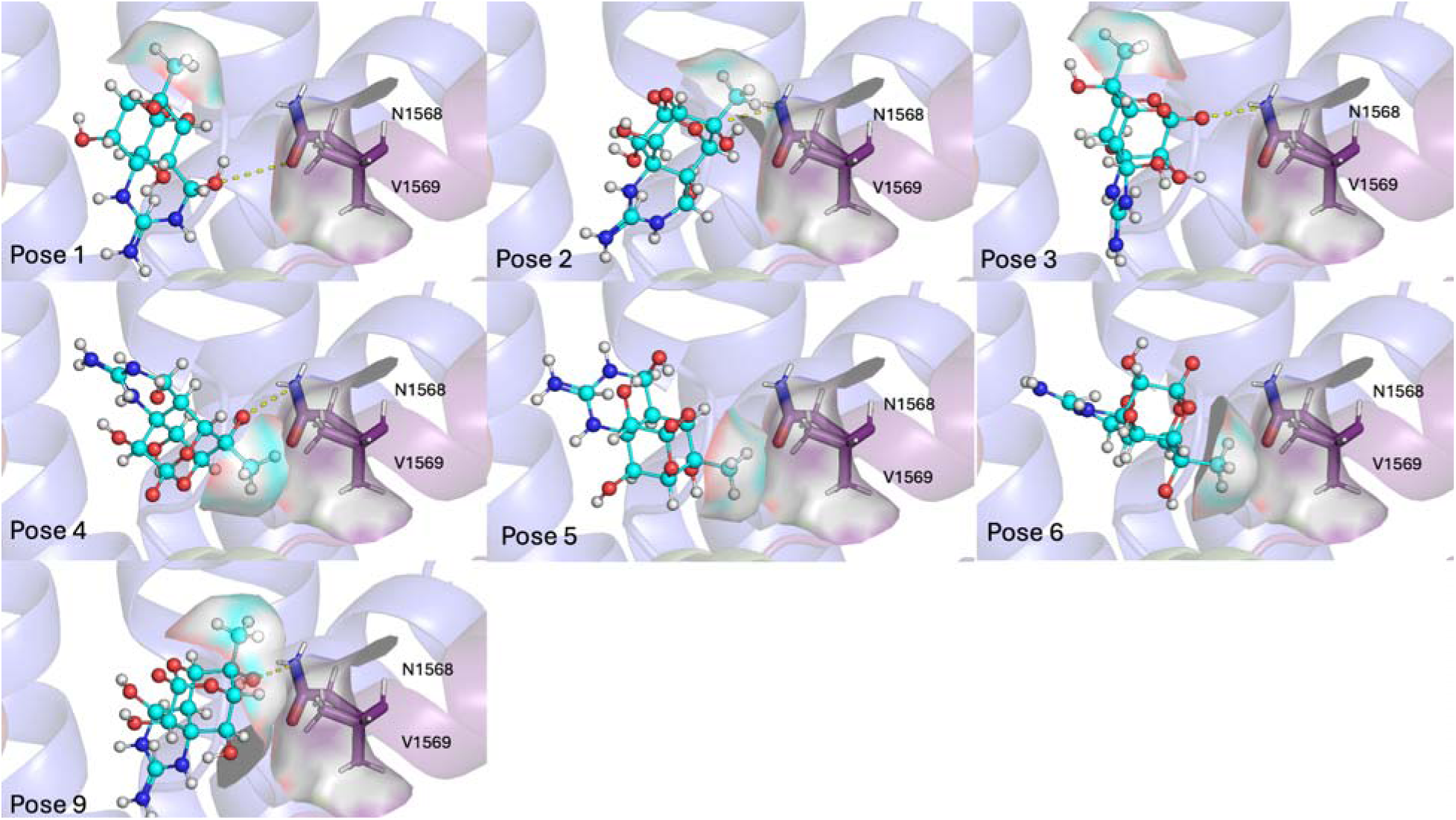
Selected poses (1, 2, 3, 4, 5, 6, 9) of the highest scoring analog, 11-deoxy-4-epi-TTX (cyan, ball and stick) in Na 1.4^LVNV^. N1568 and V1569 in Domain IV are shown as sticks with TTX-N1568 interactions shown as yellow dashes. V1569 and the C11-methyl of 11-deoxy-4-epi-TTX are shown as surface to showcase hydrophobic interactions.

## Discussion

Here, we show that Pacific newts possess a diverse arsenal of toxic weapons that may act against their garter snake predators. TTX analogs vary in richness within and among newt species, and in some cases, these analogs reach appreciable concentrations (table 2; fig. 2, 3), suggesting they may have ecological relevance in antipredator defense. As a first step toward understanding this functional relevance, we used molecular docking to assess whether how TTX variants interact with snake sodium channels compared to canonical TTX (fig. 5). We find that some of these TTX variants have the potential to block snake channels that have previously shown as virtually invulnerable to canonical TTX. Taken together, these results force a revaluation of the evolutionary tug-of-war between newts and snakes. Specifically, our analyses suggest that the diversification of TTX analogs may provide newts with an evolutionary counterattack, even against garter snake populations that have previously been characterized “winning” the arms race (Brodie et al. 2002; Hanifin et al. 2008; Feldman et al. 2010; Reimche et al. 2020).

### Analogs across *Taricha* spp

We discovered that all four Pacific newt species (*T. granulosa, T. rivularis, T. sierrae,* and *T. torosa*) possess a diversity of TTX compounds (table 2). The suite of analogs we detected includes epi-TTX, anhydro-TTX, deoxy-TTX, and several additional TTX variants. Additionally, we discovered a new TTX analog not previously reported in *Taricha* that appears to be 11-oxotetrodotoxin, a compound also found in the eastern newt, *Notophthalmus viridescens* (Yotsu-Yamashita and Mebs 2003). Previous work on TTX in *Taricha* has focused almost exclusively on the rough-skinned newt (*T. granulosa*), measuring TTX levels and a limited set of analogs such as 6-epi-TTX and 11-deoxy-TTX (Kotaki and Shimizu 1993; Hanifin et al. 1999; Kudo et al. 2020). As such, there has been almost no work identifying TTX analogs in *T. rivularis, T. sierrae,* or *T. torosa*, and our analysis shows that the diversity of TTX compounds is just as rich in these other newt species as in *T. granulosa* (table 2). These TTX analogs differ in content and abundance across the four newt species. But for all species, the most abundant compounds were most likely 4,9-anhydro-TTX (Kotaki and Shimizu 1993; Hanifin et al. 2002), 11-deoxy-TTX (Yotsu et al.1990; Kotaki and Shimizu 1993), and TTX (Buchwald et al. 1964, Wakely et al. 1966, Hanifin et al. 1999, 2002; Kudo et al. 2016), respectively. Though further analytical work will be required to elucidate the exact structures of these compounds and confirm their identity, their masses match TTX analogs previously found in *T. granulosa* and the Asian newt *Cynops ensicauda* (Yasumoto et al. 1988; Yotsu et al. 1990; Kotaki and Shimizu 1993; Hanifin et al. 2002).

### TTX Analog Potency and Sodium Channel Docking

Few of the TTX analogs have been assayed on voltage-gated sodium channels to quantify potency (i.e. channel block). Kudo and Yotsu-Yamashita (2019) showed that 6-epi-TTX and 8-epi-TTX are less potent than canonical TTX in mouse cells, consistent with the favorable interaction of the C8 hydroxyl of TTX with E801 in DII of the selectivity filter. Epimerization of this group, as in 8-epi-TTX, would potentially reduce the likelihood of this interaction. Biochemical assays would need to be performed with snake Na_V_ channels to test toxin potency given the difference in sequence. The toxicity of 4,9-anhydro-TTX has been tested in several human Na_V_ isoforms. Rosker et al. (2007) showed that TTX is 40-231 times more effective compared to 4,9-anhydro-TTX in the human Na_V_1.2-1.8 channels except for peripheral and central nerve channel Na_V_1.6, for which 4,9-anhydro-TTX was a better inhibitor. More recent work found that 4,9-anhydro-TTX also inhibits Na_V_1.1 in humans and mice (Denomme et al. 2020). However, 4,9-anhydro-TTX has been found to inhibit Na_V_1.4-1.6 isoforms, with Na_V_1.6 requiring about triple the concentration for inhibition over Na_V_1.4 and Na_V_1.5, according to NMR spectroscopy (Tsukamoto et al. 2017). Our molecular docking analyses show that this TTX analog is on par with canonical TTX with slightly less favorable scores in all Na_V_ variants except Na_V_1.4^T^ in *T. couchii* (fig. 5). Electrophysiological testing of 4,9-anhydro-TTX on garter snake Na_V_1.4 would be needed to elucidate the relative potency of this analog in the *Thamnophis* system. While 4,9-anhydro analog types are in lower relative abundance in the newts we sampled (fig. 3), possessing this analog as a potential weapon in a heterogenous landscape of snake mutations could be beneficial to some newts, especially if paired with the 11-deoxy modification. While we did not dock 4,9-anhydo-11-deoxy-TTX, it is present in other newts (table 2) (Yasumoto et al. 1988) and a plausible predicted structure for the 285.0960 experimental mass. Given the favorability of the 11-deoxy-TTX analogs across the snake Na_V_ variants (fig. 5), it is reasonable to hypothesize that 4,9-anhydro-11-deoxy-TTX could be an additional alternative effective chemical weapon.

We show that 11-deoxy-TTX variants dock more favorably than canonical TTX across the spectrum of *Thamnophis* Na_V_1.4 variants, especially in the extremely resistant *T. sirtalis* Na_V_1.4^LVNV^ channel. Coincidentally, we also found that the putative 11-deoxy-TTX analogs were among the most abundant in our *Taricha* skin secretions. However, there is evidence that 11-deoxy-TTX is less potent compared to TTX at least in Neuro-2A cells which primarily express Na_V_1.7 (Reverté et al. 2024) which is quite distinct from Na_V_1.4. Choudhary et al. (2003) mutated Na_V_1.4 D1532N (D1568N in snakes) and found a sixfold decrease in binding of 11-deoxy-TTX compared to TTX. But from TTX assays in various *Thamnophis sirtalis* Na_V_1.4 specifically, a pattern of potential synergy between the four mutations begins to emerge. Geffeney et al. (2005) tested a non-naturally occurring mutation, Na_V_1.4^LINV^ instead of Na_V_1.4^LVNV^, and showed the K_D_ decreases by half (∼12,000nM to ∼5,900nM), indicating there are interactive effects between the I1561V that does not contact TTX and the D1568N site that directly interacts with the C11 of TTX. Perhaps there are synergistic effects of the LVNV sites within themselves and the rest of the pore that are not captured solely by considering the static aspects of the point mutation. Future MD simulations may better capture these interactions to understand how the more TTX-distant I1561V residue plays a role in doubling the resistance level.

Docking does not explicitly account for desolvation, yet desolvation energetics likely impact the behaviors we observe. Desolvation refers to the energetic cost of removing water molecules that hydrate both the ligand and binding site prior to complexation. The C11 hydroxyl (-OH) group of canonical TTX forms favorable hydrogen-bonding interactions with surrounding water molecules. Consequently, when TTX binds to the channel pore it disrupts these stabilizing water interactions, imposing a relatively strong desolvation penalty. In contrast, the C11 methyl group (-CH_3_) of the 11-deoxy analogs interacts much less favorably with water, making desolvation of this analog energetically less costly. This difference becomes important in the resistant Na_V_1.4^LVNV^ pore, where the ancestral polar and negatively charged D1568 and G1569 residues (present in Na_V_1.4^+^) are replaced by the more hydrophobic N and V residues, respectively. These substitutions reduce the polarity of the pore environment and likely diminish opportunities for strong water-mediated hydrogen bonding with canonical TTX. Under these more hydrophobic pore conditions, 11-deoxy analogs may be comparatively favored because their less polar C11 methyl group incurs a smaller desolvation cost and is therefore more chemically compatibly with the more hydrophobic pore residues. Thus, 11-deoxy-TTX analogs may act as an effect weapon in the arms race because its chemical properties are particularly well suited for targeting the extremely resistant Na_V_1.4^LVNV^. Regardless of the specific biophysical interactions between the putative 11-deoxy-TTX and snake channels, this TTX analog was one of the most abundant across all newt species and may act as an effective weapon in the arms race with garter snakes.

### The Case of *T. couchii*

The Sierra garter snake (*T. couchii*) has been a puzzle. The species preys on *T. torosa* (Brodie et al. 2005) and *T. sierrae* (Wiseman and Pool 2007) and shows tremendous TTX-resistance at the southern end of its distribution grading into TTX-sensitivity moving north (Brodie et al. 2005; Feldman et al. 2009; Reimche et al. 2020). Yet the species fixed for a single Na_V_1.4 allele (Na_V_1.4^T^) which contains a DIII P-loop mutation, M1276T (Reimche et al. 2022) also seen in TTX-bearing newts, pufferfish, and octopus (Jost et al. 2008; Geffeney et al. 2019; Gendreau et al. 2021). This replacement has been verified to decrease TTX binding to the rat Na_v_1.4 roughly 15-fold (Jost et al. 2008). Yet in our docking simulations, Na_V_1.4^T^ had the greatest number of TTX analogs that score better on average than TTX (fig. 5). The M1276T replacement in *T. couchii* is engaged more often in the *T. sirtalis* Na_V_1.4^LVNV^ channel compared to the sensitive Na_V_1.4^+^ channel (fig. S4). This outcome could result from a less ideal approximation we made by introducing the mutation and not accounting for some small structural rearrangements that naturally occur in Na_V_1.4^T^. It also is possible that the Na_V_1.4^T^ channel is more TTX-sensitive than previously thought and that *T. couchii* employs an alternative mode of resistance, such as detoxification mechanisms or TTX-binding proteins (Robinson et al. 2024), which might bind multiple analogs of TTX. This mechanism would relieve channels of the selective pressure to gain and maintain multiple costly mutations to compete in the arms race.

In addition to the surprising docking results, *T. couchii* overlaps almost entirely with *T. sierrae* (Stebbins 2003; Hansen and Shedd 2025), which showed the most striking pattern of TTX analog relative abundances (fig. 3). This newt species contained the highest amounts of one of the putative 11-deoxy-TTX analogs (amu 303; compounds **6-9**), possessing more of this compound than even TTX (unlike the other *Taricha* species). Snakes that eat these newts likely experience selective pressure to maintain or revert to a hydrophilic pore to maintain resistance.

### Next Steps in the Escalation of the Arms Race

Snakes preying on newts face a narrow mutational landscape, constrained by antagonistic pleiotropy that limits the adaptive path (Feldman et al. 2012; Brodie and Brodie 2015; del Carlo et al. 2024). Overall, the sodium channels of snakes must balance the hydrogen bonding network to prevent TTX block, while still conducting sodium-specific current, and detecting proper activation voltages (del Carlo et al. 2024). For example, both D396 and E801 create increased contacts with TTX (fig. S4) but are almost certainly forbidden by selection. These sites are highly conserved in Na_v_ across metazoa as two of the four residues that form the selectivity filter (the DEKA motif), giving rise to the remarkable specificity for Na^+^ over other ions (Noda et al. 1989; Terlau et al. 1991; Dudev and Lim 2014; Zhorov 2021). When newts employ multiple TTX analogs that make slightly different contacts with the sodium channel, then the potential pool of adaptive substitutions likely shrinks further.

Because of the critical role of sodium channels, some of the Na_v_ replacements seen in garter snakes likely rescue channel function. For example, there is evidence of the hydrophobic costs of replacing the glutamate (D) with the less polar asparagine (N) (the D1568N substitution) being offset by the D1277E substitution seen in some populations of *T. atratus* that have the Na_V_1.4^EPN^ allele (del Carlo et al. 2024). By calculating interaction energies for the extremely resistant Na_V_1.4^EPN^ and Na_V_1.4^LVNV^ channels, del Carlo et al. (2024) showed that the DIII D1277E has stabilizing effects on the DIV D1568N in *T. atratus*, whereas Na_V_1.4^LVNV^ does not have any compensatory interactions to account for its hydrophobic destabilization. In fact, the N1568 shows a deleterious interaction with the DIII K1273 of the selectivity filter in both Na_V_1.4^LVNV^ and Na_V_1.4^EPN^. In our docking contacts, we see this lysine engage in a greater number of contacts in the Na_V_1.4^LVNV^ compared to Na_V_1.4^+^ that could be compensating for the increased hydrophobicity of the channel, whereas Na_V_1.4^EPN^ has fewer overall K1273 contacts compared to its moderately resistant species counterpart, Na_V_1.4^P^ (fig. S4). These findings further emphasize that the arms race is constrained by the functional demands of the Na_V_ protein itself. Selection for increased TTX resistance may therefore extend beyond TTX-interacting residues to include compensatory substitutions that preserve channel stability and normal function despite destabilizing effect of extreme resistance mutation.

There are shifts in hydrogen bonding patterns across Na_V_1.4^LVNV^ and Na_V_1.4^+^ that create a vulnerability, leaving Na_V_1.4^LVNV^ to become selectively compatible with 11-deoxy-TTX variants. These analogs favor hydrogen bond donor/acceptor interactions that rely more on amide chemistry in the N1568 than the acidic interactions of D1568 that dominate in TTX binding. While all four differences between Na_V_1.4^+^ and Na_V_1.4^LVNV^ reside in the DIV of the pore, only two of them directly interact with the TTX analogs. The TTX-contacting mutations in the resistant channel, D1568N and G1569V, make the pore more hydrophobic, properties that are more suitable for canonical TTX evasion, but favor interactions with the less polar 11-deoxy-TTX variants, whose missing C11 hydroxyl group leaves a hydrophobic terminal methyl group. Whereas some TTX interaction sites are maintained between sensitive and resistant channels, the more hydrophobic 11-deoxy-TTX analogs are more complementary to the more hydrophobic pore. As a result, 11-deoxy-TTX analogs may be a more effective weapon precisely because their chemistry is better matched to the chemistry of the extremely resistant Na_V_1.4L^VNV^ pore.

This framework generates clear predictions: (i) selection will act to increase the relative abundance of the 11-deoxy-TTX analogs in newt populations sympatric with highly resistant snakes; and (ii), snake populations exposed to 11-deoxy-TTX analogs will experience renewed selection for polar or compensatory pore substitutions. Future work should test the potency of 11-deoxy-TTX variants against both TTX-sensitive and TTX-resistant snake channels. We predict that electrophysiological assays using 11-deoxy-TTX analogs will reveal that these TTX structures block TTX-resistant Na_V_1.4 (e.g. Na_V_1.4^LVNV^) more effectively than canonical TTX. As such, the coevolutionary story surrounding newts and snakes may be more complex than previously recognized, involving multiple toxins that assault snake channels in subtle but deadly ways.

## Methods

### Newt Field Collections and Housing

We collected 40 individual Pacific newts, obtaining 10 adults of each of the four *Taricha* species: T*. granulosa, T. rivularis, T. sierrae, T. torosa* (table 1). We collected individuals by hand or hand-held net from ponds and creeks at field sites in Central and Northern California (table 1). We transported newts in coolers from field to lab (at UNR). We kept newts in a temperature-controlled room averaging 15 - 20°C where they were housed individually in 19 L glass aquaria with two PVC-pipe halves that served as both hides and haul-outs, and filled 6 - 8 cm deep with dechlorinated water. We kept newts on a 12L:12D light cycle and fed them a variety of food two or three times weekly, including frozen (thawed) brine shrimp and blood worms, as well as live earthworms. We followed protocols approved by UNR Institutional Animal Care and Use Committees (IACUC) for all care, handling, and procedures.

### Newt Toxin Sampling

Newts store defensive compounds such as TTX and TTX analogs in dermal granular glands (Mailhao-Fontana et al. 2019). We used an electrostimulation procedure similar to Cardall et al. (2004) to collect newt skin secretions from these glands. We first placed newts in a plastic tub containing a 1% solution of Tricaine methanesulfonate (MS-222) to anesthetize them. Once unconscious (unresponsive to physical touch), we rinsed newts off with tap water, patted them dry with a sterile paper towel, and placed them belly-down on a paper towel. While the newts were still sedated, we applied electric stimulation (via bifurcating cable electrodes) to their dorsal surface. We gently ran electrodes (set at 5.4 V, 10 pps, pulse duration 14 ms) down one side of the dorsum, from base of tail to tip, and then again along the other side of the body (generally 4 passes in total), gathering skin secretions with a sterile cotton swab. We then stored swabs in sterile epitubes in a −80°C freezer until later use. Following electrostimulation, we placed newts in a fresh plastic tub with a wet paper towel where they were allowed to recover for one hour. We then rinsed newts a final time in tap water and returned them to their home tanks. We observed no adverse effects or distress, maintaining newts for many months afterwards for subsequent studies.

### Newt Toxin Characterization (LC–MS Analysis)

We extracted newt skin secretions from the cotton swabs by soaking swabs in Optima-grade LC–MS methanol (Fisher Scientific, Pittsburgh, PA, USA) and sonicating swabs for 15 min in an ice bath. We filtered samples using a syringe filter (0.45 μm) into 2 mL autosampler vials. We then analyzed the skin secretion extracts with an Agilent 1290 Infinity II ultrahigh performance liquid chromatograph (UPLC; Agilent Technologies, Santa Clara, CA, USA) equipped with a binary pump, multisampler, column compartment and diode array UV/Vis detector, coupled to an Agilent 6560 Ion Mobility Quadrupole Time-of-Flight (IM-QTOF) mass spectrometer via a Jet Stream electrospray ionization source (ESI-TOF; gas temperature: 300 °C, flow: 11 L/m; nebulizer pressure: 35 psig; sheath gas temp: 300 °C; sheath gas flow: 11 L/m; VCap: 3500 V; nozzle voltage: 500V; fragmentor: 350 V; skimmer: 65 V; octopole: 750 V). We analyzed samples in low-mass (100-1700 m/z) TOF mode at a rate of 1 spectrum/sec (8131 transients/spectrum). We co-injected extracts (1.00 μL) with nicotine internal standard (1.00 uL, 10.0 μM in MeOH) and eluted at 0.400 mL/min through a Kinetex Polar C18 column (Phenomenex: 2.1 x 150 mm, 2.6 μ, 100 Å) at 40 °C. The linear binary gradient is comprised of buffers A (water containing 10 mM ammonium acetate) and B (acetonitrile containing 1 % water) changing over 20 minutes accordingly: hold 0-4 min 0% B, ramp to 30% B at 12 min, ramp to 100% B at 15 min, 15-17 min hold at 100% B, 17.1-20 min hold at 0% B. We extracted peaks, corrected retention time, aligned and grouped using recursive small molecule analysis in Agilent Profinder v10.0 with a minimum height cut-off of 20,000 counts. We imported data into Agilent Mass Profiler Professional v15.1 where peak areas were normalized to nicotine internal standard. We used the resulting entity list (236 m/z, rt bins) for statistical analyses. We deconvoluted these 236 LC–MS entities by grouping entities within 0.1 min retention time of one another and having a Pearson correlation ≥ 0.7 yielding 161 putative compounds before normalizing to secretion dry weight and extraction volume. We also analyzed a blank extraction of a cotton swab without secretion to excise any peaks that might have resulted from the swab. We pooled newt extracts by species and analyzed in Q-TOF mode using iterative data-dependent acquisition (collision energy = 20, 40 V; precursor threshold = 10k counts).

### Newt TTX Analog Structures

Of the 161 unique compounds detected in extract of newt skin secretions, we identified nine unique TTX analogs (see Results; table 2). We identified compounds as TTX analogs based on molecular mass (and retention time) in comparison to previously published literature on TTX analogs in newts (Yostu and Yasumoto 1990; Yotsu et al. 1990; Kotaki & Shimizu 1993; Hanifin et al. 1999; Yostu-Yamashita 2001; Hanifin et al. 2002; Yotsu-Yamashita and Mebs 2003; Hanifin 2010; Kudo et al. 2014; Silva et al. 2019; Kudo et al. 2020; Hanifin et al. 2022). However, without known standards (only purified TTX was available to us) we were unable to fully identify all possible isobaric analogs. We limited our selection to analogs previously isolated and characterized from newts (family Salamandridae) in the literature, and excluded synthetic analogs proposed in the TTX biosynthetic pathway. We chose TTX analogs having a theoretical mass within 10 ppm of the experimentally determined neutral molecular masses. This procedure resulted in 14 possible known TTX analog structures from literature (fig. 2 and table 2). We then selected a subset of those analogs between 301 and 319 amu (**1-4**, **6-9**, **12**, **13**) to prepare for molecular docking (fig. 3). To determine the influence of protonation state of the C-10 oxygen in hemilactal form, we also included a form of canonical TTX with this C-10 oxygen protonated. Given the oxyanionic form of TTX scored more favorably on average in all Na_V_ variants compared to the protonated hydroxyl (fig. S2), we opted to dock all the hemilactal TTX analogs in the oxyanionic form (**1-4**, **6**, **7**). We obtained available structures from PubChem and generated unavailable TTX analogs using the structure editing features in Avogadro1.2.0 (Hanwell et al. 2012). We imported structures into AutoDock Tools (ADT) (MGLTools1.5.6) (Morris et al. 2009 to assign charges and to generate torsion trees, with manual adjustments to restrain non-rotatable bonds that were not automatically detected. We generated PDBQT files and then corrected charges using the semiempirical AM1-BCC charge calculations in Antechamber from AmberTools25 (Case et al. 2023).

### Newt TTX Analog Variation

We conducted two statistical analyses to quantify variation in TTX analogs within and among newt species. We performed all statistical analyses and visualizations with R studio v2026.01.1+403 (R Core Development Team 2026) and the R package gplots v3.3.0 (Warnes et al. 2025). We log_2_ transformed all relative compound abundance data to normalize the data. To assess how TTX analogs differ among newt species, we conducted ANOVA examining the effect of newt species and TTX analog (with an interaction) on relative analog abundance, using a pairwise Tukey’s post hoc analysis to identify significantly different contrasts between species. Lastly, we used Simpson’s diversity index to calculate TTX analog diversity for each species using vegan v2.7-3 in R.

### Snake Skeletal Muscle Sodium Channel (Na_V_1.4) Structures

We included the sodium channel variants found in wild populations of all three species of *Thamnophis* (*T. atratus*, *T. couchii*, *T. sirtalis*) that prey on *Taricha* as targets for molecular docking. These species possess at least seven distinct skeletal muscle sodium channel (Na_V_1.4) variants as follows (fig. 1): Na_V_1.4^+^ the ancestral TTX sensitive Na_V_1.4 that contains no pore loop (P-loop) mutations and is seen in nearly all garter snake species, including populations of *T. atratus* and *T. sirtalis* (Geffeney et al. 2005; Feldman et al. 2009; Hague et al., 2017); Na_V_1.4^P^ containing a single replacement in the DIII P-loop of *T. atratus* associated with low levels of TTX resistance (Feldman et al. 2010; Moniz et al. 2022); Na_V_1.4^EPN^ with three substitutions in the pore of *T. atratus* conferring extreme TTX resistance (Feldman et al. 2010; Moniz et al. 2022; del Carlo et al. 2024); Na_V_1.4^T^ containing a single replacement in the DIII P-loop of *T. couchii* associated with high to variable levels of TTX resistance (Feldman et al. 2009; Reimche et al. 2022); Na_V_1.4^V^ containing a single mutation in the DIV P-loop of *T. sirtalis* conferring low levels of TTX resistance and largely seen in Oregon populations (Geffeney et al. 2005; Hague et al., 2017, 2020); Na_V_1.4^VA^ with two mutations in the DIV P-loop of *T. sirtalis* conferring high levels of TTX resistance, found in some Oregon and Washington populations (Geffeney et al. 2005; Hague et al., 2017, 2020); and Na_V_1.4^LVNV^ with four replacements in the DIV P-loop of some California populations of *T. sirtalis* that provides extreme TTX resistance (Geffeney et al. 2005; Feldman et al. 2009, 2010; Hague et al., 2017, 2020; Moniz et al. 2022; del Carlo et al. 2024). Though nearly all of these sodium channel variants have been constructed and expressed *ex vivo* to verify the effects of pore loop replacements on TTX binding (Geffeney et al. 2005; Jost et al. 2008; del Carlo et al. 2024), none of these channels have actually been experimentally resolved (i.e. there is no structure of the protein available to provide an empirical map 3D map of the channel). We therefore needed to generate models of each of the seven snake sodium channel variants. An experimental structure of a cockroach Na_V_PaS channel (PDB: 6A95) has been resolved via cryo-electron microscopy with TTX bound (Shen et al. 2018) and a Japanese pufferfish (*Takifugu rubripes*) Na_V_1.4a channel has been modeled by AlphaFold (Jumper et al. 2021). These two structures provide the basis for the *Thamnophis* Na_V_1.4 we examined. To build a model of the extremely resistant Na_V_1.4^LVNV^ found in some populations of *T. sirtalis* (GenBank: AAW68224.1), we performed comparative modeling using Robetta (Raman et al. 2009; Song et al. 2013; Baek et al. 2021). The template we used for this modeling was the Japanese pufferfish (*Takifugu rubripes*) Na_V_1.4a channel (UNIPROT: Q2XVR7) structure predicted by AlphaFold (Jumper et al. 2021). This channel is also TTX-resistant (Venkatesh et al. 2005, Jost et al. 2008), is very similar in length to the snake channel (1875 and 1892 amino acids for *T. sirtalis* and *T. rupripes*, respectively) and has 63.9% sequence identity and 82.6% similarity using the BLOSUM45 matrix. We aligned all five Robetta output models and selected one that had a C-terminal region that localized close to the rest of the protein and could be reasonably positioned close to a phospholipid membrane. Three of the Robetta models had extended, disordered C-terminal regions (fig. S1) and were discarded as unlikely to be reliable predictions. The remaining two models (fig. S1) were verified via structural assessment metrics in SWISS-MODEL (Waterhouse et al. 2024). The best model exhibited reasonable QMEANBrane (Studer et al. 2014) scores with the transmembrane region having high quality scores throughout (fig. S1). Some regions outside of the membrane region have lower quality scores, which is expected given the disordered nature of these regions. Ramachandran plots showed that nearly all residues adopted φ and ψ angles in allowed or generously allowed regions, with few outliers (fig. S1).

As a first step in predicting the structures of six other snake channels, we first generated a structure of the ancestral TTX-sensitive snake channel (Na_V_1.4^+^), using the Na_V_1.4 sequence from a TTX sensitive *T. sirtalis* (GenBank: AYG96483) modeled from our Robetta model of Na_V_1.4^LVNV^. To ensure that the conformation of the resulting snake channels would be conducive to docking TTX, we performed a short molecular dynamics (MD) simulation to equilibrate the Na_V_1.4^+^ structure with TTX present. Doing so ensured that the pore was in a configuration with a binding pocket that would be amenable to docking TTX. Performing these MD simulations required that the protein be placed in a hydrated palmitoyloleoylphosphatidylcholine (POPC) membrane. We used CHARMM-GUI v3.7 Membrane Builder (Jo et al. 2007; Jo et al. 2008; Brooks et al. 2009; Wu et al 2014; Lee et al. 2016; Lee et al. 2018) to build a Na_V_-POPC membrane system, specifying disulfide bonds analogous to those in the sodium channel of the American cockroach (*Periplaneta americana*), Na_V_PaS (Shen et al. 2017) (Na_V_1.4^+^: 278-359, 350-365, 806-815, 1215-1235, 1582-1597), and the 14 mutations (K42E, L184F, I332F, R542D, V600A, S714F, K865R, S899N, L1556I, V1561I, N1568D, V1569G, P1792Q, G1798E) to convert the Robetta model of Na_V_1.4^LVNV^ to the sensitive Na_V_1.4^+^ channel. The system contained 583 lipids, 300 in the upper leaflet and 283 in the lower leaflet. Water was added to a thickness of 30 Å and was represented by the CHARMM-modified TIP3P model (Jorgensen et al. 1983, Durell et al. 1994, Neria et al. 1996). NaCl was added to reach a concentration of 150 mM using standard CHARMM ion parameters (Beglov and Roux 1994). We used the CHARMM36m force field (Mackerell et al. 1998; Mackerell et al. 2004; Best et al. 2012; Huan et al. 2017) to represent the proteins and the CHARMM36 force field (Klauda et al. 2010; Klauda et al. 2012) to represent the lipids.

We generated the topology for TTX using CGenFF v4.6 (Vanommeslaeghe et al. 2012) and refined by symmetrizing charges on equivalent groups and reassigning charges to the guanidinium moiety based on the standard CHARMM methylguanidinium model compound. We refined dihedral parameters with large parameter penalties by comparing the quantum mechanically optimized geometry (MP2/6-31+G*) to that of the molecular mechanics model. Given that there were no rotatable bonds in TTX that required refinement, we could not perform one-dimensional energy scans, so we considered agreement with the optimal geometry sufficient. We then minimized the Na_V_1.4-TTX system via 5000 steps of steepest descent via the CHARMM program (Hwang et al. 2024) and then equilibrated in OpenMM 7.5.1 (Eastman et al. 2017) for 50 ns following the CHARMM-GUI Membrane Builder protocol. We used the protein structure at the end of the equilibration as the structure input for making all *Thamnophis* Na_V_ channels used in molecular docking. We removed TTX, water, ions, and phospholipids prior to docking.

We used the equilibrated model of the *T. sirtalis* Na_V_1.4^+^ channel as the base structure for the remaining snake channel variants (fig. 1). The modeled structure was uploaded to CHARMM-GUI to specify disulfide bonds and introduce mutations into the pore of Domains III and IV for each corresponding Na_V_ channel (fig. 1). Each protein was minimized with 100 steps of steepest descent to relax any poor contacts induced by the mutations. Structures were edited in ADT to merge non-polar hydrogen atoms, assign AD4 atom types, and compute Gasteiger charges. The net protein charge was compared and verified against the charges calculated by CHARMM based on standard residue definitions of the CHARMM36m force field (Huang et al. 2017). Protonation states of titratable residues were verified by comparison against the topology generated by CHARMM-GUI, ensuring consistency of the minimized structure with the adjusted structure prepared by ADT. In some cases, arginine charges were not assigned correctly by ADT and were manually adjusted to the standard value in the PDBQT files.

### Molecular Docking

We performed docking using AutoDock Vina v1.1.2 (Trott and Olson 2010). We docked 10 of the 14 TTX analogs (compounds **1-4**, **6-9**, **12**, **13**; fig. 2) into each of the seven snake Na_V_1.4 variants and Na_V_PaS, generating nine ligand poses (the default setting in Vina). We used ADT to determine the search space parameters with the box centered on TTX from the previously equilibrated TTX-sensitive *Thamnophis* channel (1.4^+^). We centered the box at (0.283, 0.051, 27.306 Å) and included 30 x 32 x 18 grid points with 1.0-Å spacing to cover the pore loops. We structurally aligned all the sodium channels prior to molecular docking and used the same parameters across every snake channel. Molecular docking returns a value known as a docking score from an empirical scoring function rather than a direct estimate of binding affinity (K_D_). Important factors such as desolvation penalties and entropic contributions of the protein and ligand are not considered. Therefore, the docking outcomes are semi-quantitative approximations of affinity and may not be directly comparable to measures of phenotypic resistance to TTX in garter snakes obtained through whole-animal assay (e.g. Brodie et al. 2002; Reimche et al. 2020). Nevertheless, this approach can reveal overall trends in interactions between TTX analogs and Na_V_ variants that can be further investigated. Using molecular docking to explore Na_V_-TTX interactions is ideal to address our hypotheses because it is a relatively computationally inexpensive approach, it provides atomic level understanding for an ecological interaction, and importantly, most TTX analogs are not commercially available for comparable techniques that would provide this atomic level insight.

### Redocking to the Cockroach (Na_V_PaS) Channel

To validate our docking approach, we included the cockroach sodium channel, Na_V_PaS (PDB ID: 6A95) (Shen et al. 2018), to perform a redocking protocol because this structure was originally resolved with TTX bound. We used MODELLER v10.5 (Fiser et al. 2000; Webb and Sali 2016) to construct two missing intracellular loops (residues 436-508 and 745-839) and proceeded with the best scoring structure. We then subjected this modeled structure to the same protocol described above (for Na_V_1.4^+^). Using CHARMM-GUI, we specified disulfide bonds based on cryo-EM data (288-337, 328-343, 709-717, 1011-1030, and 1368-1381) and we inserted the protein into a POPC membrane. The system contained 670 lipids, 340 in the upper leaflet and 330 in the lower leaflet. Following minimization and equilibration as described above, we removed water, lipids, and ions and used the equilibrated protein structure at the input for TTX docking.

### Docking Analyses

We used the docking scores generated by Vina to calculate the average docking score across the nine poses for each TTX analog for each snake Na_V_1.4 variant. We visualized each docking relative to the score for canonical TTX in the sensitive snake channel Na_V_1.4^+^ (fig. 5). We then took the best-scoring TTX analogs (relative to canonical TTX) in the extremely resistant *T. sirtalis* Na_V_1.4^LVNV^ channel and investigated the differences in amino acid contacts using interaction fingerprints generated by Schrodinger Maestro14.3 (Duan et al. (fig. 6, 7). Doing so allowed us to determine which types of interactions were responsible for the more favorable average docking scores. Additionally, we investigated overall contacts between all channels and TTX analogs using interaction fingerprints generated by Schrodinger Maestro14.3 (fig. S4) (Duan et al. 2010; Sastry et al. 2010). The overall contacts include the following possible types of interactions: aromatic, hydrophobic, hydrogen bond donor, hydrogen bond acceptor, polar, charged, sidechain, and backbone.

## Supporting information

Raw abundance data

## Supplementary material

Supplementary material is available.

## Acknowledgments

We thank California Department of Fish and Wildlife (CAF&W) for permits to C.R.F., and UNR IACUC for approval of live animal protocols (#21-01-115-1) to C.R.F. and K.E.R. We are grateful to M. Rawlins (Sonoma State University) for access to sampling sites. We thank J. Wilcox and E. Taylor for assisting with animal collection, as well as UNR Office of Animal Resources for live animal support. We thank M. LoPresti for Python scripts used for parsing interaction fingerprints generated by Schrodinger. We appreciate useful feedback on this manuscript from the UNR Evol Doers, especially S. Babish, D. Alvarez-Ponce, K. Wojczulanis-Jakubas, and E. McMullen.

## Data availability

Raw data from LC–MS runs will be uploaded to a public database. Computational models rely on these publicly available sources: Protein Data Bank (PDB 6A95), GenBank (AAW68224.1 and AYG96483), and UNIPROT (Q2XVR7).

