## Supporting tables and figures for "Pick Your Poison: Tetrodotoxin Variants Give Pacific Newts a Potential Leg Up in the Coevolutionary Arms Race with Resistant Garter Snake Predators"

**Table S1.** Raw docking scores for all nine poses of each of the 14 total docked ligands (including protonated TTX) in each of the seven $Na_V$ channels and the cockroach model ($Na_V$PaS). Mean score across all nine poses given in the last column.

| Protein Variant | TTX Analog | Pose 1 | Pose 2 | Pose 3 | Pose 4 | Pose 5 | Pose 6 | Pose 7 | Pose 8 | Pose 9 | Mean |
|---|---|---|---|---|---|---|---|---|---|---|---|
| Nav1.4$^{LVNV}$ | 8-epi-TTX | -7.6 | -7.2 | -7.2 | -7 | -7 | -6.7 | -6.7 | -6.6 | -6.6 | -6.96 |
| Nav1.4$^{LVNV}$ | 11-deoxy-4-epi-TTX | -8.4 | -7.7 | -7.5 | -7.5 | -7.5 | -7.3 | -7.1 | -7.1 | -7 | -7.46 |
| Nav1.4$^{LVNV}$ | 4,9-anhydro-TTX | -7.6 | -7.3 | -7.3 | -7.2 | -7.2 | -7.1 | -6.9 | -6.8 | -6.6 | -7.11 |
| Nav1.4$^{LVNV}$ | TTX | -8 | -7.4 | -7.3 | -7.2 | -7.1 | -7 | -6.9 | -6.9 | -6.5 | -7.14 |
| Nav1.4$^{LVNV}$ | TTX (protonated) | -7.9 | -7.3 | -7.2 | -7.1 | -7 | -6.9 | -6.9 | -6.8 | -6.8 | -7.10 |
| Nav1.4$^{LVNV}$ | 6-epi-TTX | -7.9 | -7.3 | -6.9 | -6.8 | -6.8 | -6.8 | -6.8 | -6.7 | -6.7 | -6.97 |
| Nav1.4$^{LVNV}$ | 11-deoxy-lactone-TTX | -8 | -7.5 | -7.1 | -7.1 | -6.8 | -6.7 | -6.7 | -6.6 | -6.6 | -7.01 |
| Nav1.4$^{LVNV}$ | 1-hydroxy-8-epi-5,11-dideoxy-TTX | -7 | -7 | -7 | -6.7 | -6.6 | -6.6 | -6.6 | -6.4 | -6.4 | -6.70 |
| Nav1.4$^{LVNV}$ | 4,9-anhydro-6-epi-TTX | -7.6 | -7.3 | -7.3 | -7.1 | -6.9 | -6.8 | -6.8 | -6.8 | -6.7 | -7.03 |
| Nav1.4$^{LVNV}$ | 11-deoxy-TTX | -8.3 | -7.5 | -7.3 | -7.3 | -7.2 | -7.2 | -7.2 | -6.9 | -6.9 | -7.31 |
| Nav1.4$^{LVNV}$ | 4-epi-TTX | -7.7 | -7.4 | -7.3 | -7.3 | -7 | -6.9 | -6.9 | -6.8 | -6.8 | -7.12 |
| Nav1.4$^{V}$ | 8-epi-TTX | -7.2 | -7.2 | -7.1 | -7 | -6.9 | -6.8 | -6.8 | -6.5 | -6.4 | -6.88 |
| Nav1.4$^{V}$ | 11-deoxy-4-epi-TTX | -8 | -7.4 | -7.3 | -7.2 | -7.1 | -7.1 | -7.1 | -7.1 | -6.9 | -7.24 |
| Nav1.4$^{V}$ | 4,9-anhydro-TTX | -7.6 | -7.3 | -7.2 | -7.2 | -7 | -6.9 | -6.9 | -6.8 | -6.8 | -7.08 |
| Nav1.4$^{V}$ | TTX | -7.7 | -7.3 | -7.2 | -7.1 | -6.9 | -6.9 | -6.8 | -6.8 | -6.7 | -7.04 |
| Nav1.4$^{V}$ | TTX (protonated) | -7.6 | -7.2 | -7.1 | -7 | -7 | -6.9 | -6.8 | -6.7 | -6.6 | -6.99 |
| Nav1.4$^{V}$ | 6-epi-TTX | -7.7 | -7.4 | -7 | -6.9 | -6.7 | -6.6 | -6.4 | -6.3 | -6.3 | -6.81 |
| Nav1.4$^{V}$ | 11-deoxy-lactone-TTX | -7.7 | -7.2 | -7.1 | -6.9 | -6.8 | -6.7 | -6.7 | -6.6 | -6.6 | -6.92 |
| Nav1.4$^{V}$ | 1-hydroxy-8-epi-5,11-dideoxy-TTX | -7 | -6.9 | -6.6 | -6.6 | -6.5 | -6.4 | -6.3 | -6.1 | -6.1 | -6.50 |
| Nav1.4$^{V}$ | 4,9-anhydro-6-epi-TTX | -7.7 | -7.6 | -7 | -6.9 | -6.7 | -6.7 | -6.7 | -6.7 | -6.6 | -6.96 |
| Nav1.4$^{V}$ | 11-deoxy-TTX | -8 | -7.4 | -7.3 | -7.2 | -7.1 | -7 | -6.9 | -6.7 | -6.6 | -7.13 |
| Nav1.4$^{V}$ | 4-epi-TTX | -7.4 | -7.4 | -7.4 | -7.3 | -6.9 | -6.8 | -6.7 | -6.7 | -6.7 | -7.03 |
| Nav1.4$^{VA}$ | 8-epi-TTX | -7.3 | -7.3 | -7.1 | -7.1 | -6.9 | -6.9 | -6.8 | -6.7 | -6.7 | -6.98 |
| Nav1.4$^{VA}$ | 11-deoxy-4-epi-TTX | -7.9 | -7.4 | -7.2 | -7.2 | -7.2 | -7.1 | -7.1 | -7.1 | -6.9 | -7.23 |
| Nav1.4$^{VA}$ | 4,9-anhydro-TTX | -7.5 | -7.3 | -7.2 | -7.2 | -7.2 | -7.1 | -6.9 | -6.9 | -6.8 | -7.12 |
| Nav1.4$^{VA}$ | TTX | -7.6 | -7.2 | -7.2 | -7.1 | -7 | -7 | -6.9 | -6.9 | -6.9 | -7.09 |
| Nav1.4$^{VA}$ | TTX (protonated) | -7.4 | -7.3 | -7.1 | -7 | -6.9 | -6.8 | -6.8 | -6.7 | -6.5 | -6.94 |
| Nav1.4$^{VA}$ | 6-epi-TTX | -7.6 | -7.4 | -7.1 | -7 | -6.9 | -6.8 | -6.7 | -6.5 | -6.4 | -6.93 |
| Nav1.4$^{VA}$ | 11-deoxy-lactone-TTX | -7.5 | -7.4 | -7.2 | -6.8 | -6.7 | -6.6 | -6.6 | -6.6 | -6.5 | -6.88 |
| Nav1.4$^{VA}$ | 1-hydroxy-8-epi-5,11-dideoxy-TTX | -7.1 | -6.9 | -6.7 | -6.6 | -6.6 | -6.5 | -6.5 | -6.4 | -6.1 | -6.60 |
| Nav1.4$^{VA}$ | 4,9-anhydro-6-epi-TTX | -7.7 | -7.6 | -6.9 | -6.8 | -6.7 | -6.7 | -6.7 | -6.4 | -6.2 | -6.86 |
| Nav1.4$^{VA}$ | 11-deoxy-TTX | -7.9 | -7.5 | -7.3 | -7.1 | -7.1 | -7 | -7 | -6.9 | -6.9 | -7.19 |
| Nav1.4$^{VA}$ | 4-epi-TTX | -7.4 | -7.4 | -7.3 | -7.3 | -7 | -6.9 | -6.8 | -6.7 | -6.7 | -7.06 |
| Nav1.4 | 8-epi-TTX | -7.4 | -7.3 | -7.2 | -7.2 | -7 | -6.9 | -6.9 | -6.8 | -6.8 | -7.06 |
| Nav1.4 | 11-deoxy-4-epi-TTX | -8.3 | -7.4 | -7.4 | -7.2 | -7.2 | -7.1 | -7.1 | -6.9 | -6.9 | -7.28 |

| | | | | | | | | | | | |
|---|---|---|---|---|---|---|---|---|---|---|---|
| Nav1.4 | 4,9-anhydro-TTX | -7.6 | -7.3 | -7.2 | -7.2 | -7.1 | -7.1 | -7 | -7 | -6.9 | -7.16 |
| Nav1.4 | TTX | -7.9 | -7.2 | -7.2 | -7.2 | -7.2 | -7.2 | -7.1 | -6.9 | -6.9 | -7.20 |
| Nav1.4 | TTX (protonated) | -7.7 | -7.3 | -7.2 | -7.2 | -7.2 | -7.1 | -7 | -6.8 | -6.8 | -7.14 |
| Nav1.4 | 6-epi-TTX | -7.8 | -7.5 | -7.1 | -6.9 | -6.7 | -6.7 | -6.6 | -6.6 | -6.5 | -6.93 |
| Nav1.4 | 11-deoxy-lactone-TTX | -7.9 | -7.4 | -7 | -7 | -6.9 | -6.9 | -6.6 | -6.6 | -6.6 | -6.99 |
| Nav1.4 | 1-hydroxy-8-epi-5,11-dideoxy-TTX | -6.9 | -6.8 | -6.8 | -6.8 | -6.7 | -6.7 | -6.6 | -6.5 | -6.4 | -6.69 |
| Nav1.4 | 4,9-anhydro-6-epi-TTX | -7.8 | -7.5 | -7 | -6.9 | -6.9 | -6.8 | -6.7 | -6.7 | -6.6 | -6.99 |
| Nav1.4 | 11-deoxy-TTX | -8.2 | -7.4 | -7.3 | -7.2 | -7.2 | -7.1 | -7.1 | -7 | -6.9 | -7.27 |
| Nav1.4 | 4-epi-TTX | -7.6 | -7.5 | -7.2 | -7.2 | -7.2 | -6.9 | -6.9 | -6.8 | -6.7 | -7.11 |
| Nav1.4$^{EPN}$ | 8-epi-TTX | -7.4 | -7.3 | -7.2 | -7.2 | -6.9 | -6.8 | -6.8 | -6.6 | -6.6 | -6.98 |
| Nav1.4$^{EPN}$ | 11-deoxy-4-epi-TTX | -8 | -7.4 | -7.4 | -7.3 | -7.3 | -7.1 | -7.1 | -7.1 | -6.8 | -7.28 |
| Nav1.4$^{EPN}$ | 4,9-anhydro-TTX | -7.5 | -7.3 | -7.3 | -7.2 | -7.2 | -7 | -7 | -6.9 | -6.8 | -7.13 |
| Nav1.4$^{EPN}$ | TTX | -7.7 | -7.3 | -7.2 | -7.2 | -7.1 | -7.1 | -7 | -6.9 | -6.8 | -7.14 |
| Nav1.4$^{EPN}$ | TTX (protonated) | -7.6 | -7.3 | -7.2 | -7.2 | -7.1 | -7.1 | -6.9 | -6.8 | -6.8 | -7.11 |
| Nav1.4$^{EPN}$ | 6-epi-TTX | -7.7 | -7.4 | -7.1 | -7 | -7 | -6.8 | -6.7 | -6.7 | -6.4 | -6.98 |
| Nav1.4$^{EPN}$ | 11-deoxy-lactone-TTX | -7.7 | -7.2 | -7.1 | -7.1 | -7.1 | -6.7 | -6.6 | -6.4 | -6.3 | -6.91 |
| Nav1.4$^{EPN}$ | 1-hydroxy-8-epi-5,11-dideoxy-TTX | -7 | -7 | -6.8 | -6.7 | -6.7 | -6.6 | -6.6 | -6.4 | -6.4 | -6.69 |
| Nav1.4$^{EPN}$ | 4,9-anhydro-6-epi-TTX | -7.7 | -7.6 | -6.9 | -6.8 | -6.7 | -6.7 | -6.7 | -6.6 | -6.4 | -6.90 |
| Nav1.4$^{EPN}$ | 11-deoxy-TTX | -8.1 | -7.5 | -7.3 | -7.3 | -7.2 | -7.2 | -7.1 | -6.9 | -6.7 | -7.26 |
| Nav1.4$^{EPN}$ | 4-epi-TTX | -7.5 | -7.5 | -7.4 | -7.3 | -7.1 | -6.9 | -6.7 | -6.7 | -6.6 | -7.08 |
| Nav1.4$^{P}$ | 8-epi-TTX | -7.3 | -7.2 | -7.1 | -7.1 | -7.1 | -6.9 | -6.8 | -6.6 | -6.5 | -6.96 |
| Nav1.4$^{P}$ | 11-deoxy-4-epi-TTX | -8 | -7.4 | -7.2 | -7.1 | -7.1 | -7.1 | -7.1 | -7 | -6.9 | -7.21 |
| Nav1.4$^{P}$ | 4,9-anhydro-TTX | -7.6 | -7.3 | -7.2 | -7.2 | -7.2 | -7.2 | -7 | -6.8 | -6.8 | -7.14 |
| Nav1.4$^{P}$ | TTX | -7.7 | -7.3 | -7.2 | -7.2 | -7.1 | -7 | -6.9 | -6.9 | -6.8 | -7.12 |
| Nav1.4$^{P}$ | TTX (protonated) | -7.7 | -7.3 | -7.2 | -7.2 | -7 | -6.9 | -6.8 | -6.8 | -6.7 | -7.07 |
| Nav1.4$^{P}$ | 6-epi-TTX | -7.7 | -7.4 | -7.1 | -7 | -6.9 | -6.7 | -6.6 | -6.6 | -6.6 | -6.96 |
| Nav1.4$^{P}$ | 11-deoxy-lactone-TTX | -7.7 | -7.2 | -7.1 | -6.9 | -6.7 | -6.6 | -6.5 | -6.5 | -6.5 | -6.86 |
| Nav1.4$^{P}$ | 1-hydroxy-8-epi-5,11-dideoxy-TTX | -7 | -6.9 | -6.7 | -6.6 | -6.5 | -6.4 | -6.4 | -6.3 | -6.2 | -6.56 |
| Nav1.4$^{P}$ | 4,9-anhydro-6-epi-TTX | -7.7 | -7.6 | -6.9 | -6.8 | -6.7 | -6.7 | -6.7 | -6.7 | -6.6 | -6.93 |
| Nav1.4$^{P}$ | 11-deoxy-TTX | -8.1 | -7.3 | -7.2 | -7.1 | -7 | -6.9 | -6.9 | -6.9 | -6.7 | -7.12 |
| Nav1.4$^{P}$ | 4-epi-TTX | -7.4 | -7.4 | -7.4 | -7.3 | -7 | -6.9 | -6.9 | -6.9 | -6.8 | -7.11 |
| Nav1.4$^{T}$ | 8-epi-TTX | -7.5 | -7.4 | -7.3 | -7.2 | -7.1 | -7.1 | -7 | -7 | -6.9 | -7.17 |
| Nav1.4$^{T}$ | 11-deoxy-4-epi-TTX | -8.4 | -7.5 | -7.4 | -7.3 | -7.2 | -7.1 | -7.1 | -7 | -7 | -7.33 |
| Nav1.4$^{T}$ | 4,9-anhydro-TTX | -7.6 | -7.6 | -7.5 | -7.2 | -7.2 | -7.1 | -7.1 | -6.9 | -6.8 | -7.22 |
| Nav1.4$^{T}$ | TTX | -7.8 | -7.3 | -7.3 | -7.2 | -7.1 | -7.1 | -7 | -7 | -7 | -7.20 |
| Nav1.4$^{T}$ | TTX (protonated) | -7.8 | -7.3 | -7.2 | -7.2 | -7.1 | -7.1 | -7.1 | -7 | -6.8 | -7.18 |
| Nav1.4$^{T}$ | 6-epi-TTX | -8 | -7.6 | -7.3 | -7.2 | -7.1 | -6.9 | -6.8 | -6.8 | -6.7 | -7.16 |
| Nav1.4$^{T}$ | 11-deoxy-lactone-TTX | -8.2 | -7.4 | -7.1 | -7.1 | -7 | -6.8 | -6.6 | -6.6 | -6.5 | -7.03 |
| Nav1.4$^{T}$ | 1-hydroxy-8-epi-5,11-dideoxy-TTX | -7 | -7 | -6.7 | -6.6 | -6.6 | -6.6 | -6.5 | -6.4 | -6.4 | -6.64 |
| Nav1.4$^{T}$ | 4,9-anhydro-6-epi-TTX | -7.8 | -7.6 | -7.5 | -7.1 | -7 | -7 | -7 | -6.8 | -6.7 | -7.17 |
| Nav1.4$^{T}$ | 11-deoxy-TTX | -8.3 | -7.5 | -7.4 | -7.3 | -7.2 | -7.1 | -7.1 | -7 | -7 | -7.32 |
| Nav1.4$^{T}$ | 4-epi-TTX | -7.9 | -7.5 | -7.4 | -7.4 | -7.3 | -7.3 | -7.2 | -7.1 | -6.9 | -7.33 |

| | | | | | | | | | | | |
|---|---|---|---|---|---|---|---|---|---|---|---|
| NavPaS | 8-epi-TTX | -8 | -7 | -6.8 | -6.8 | -6.6 | -6.5 | -6.4 | -6.3 | -6.3 | -6.74 |
| NavPaS | 11-deoxy-4-epi-TTX | -8 | -6.8 | -6.8 | -6.7 | -6.7 | -6.5 | -6.5 | -6.2 | -6.1 | -6.70 |
| NavPaS | 4,9-anhydro-TTX | -7.6 | -7.2 | -6.6 | -6.6 | -6.4 | -6.3 | -6.2 | -6 | -6 | -6.54 |
| NavPaS | TTX | -8.2 | -6.9 | -6.7 | -6.7 | -6.5 | -6.4 | -6.4 | -6.3 | -6.1 | -6.69 |
| NavPaS | TTX (protonated) | -8.1 | -6.6 | -6.5 | -6.4 | -6.3 | -6.3 | -6.3 | -6.3 | -6.3 | -6.57 |
| NavPaS | 6-epi-TTX | -8.2 | -7.1 | -6.7 | -6.6 | -6.3 | -6.3 | -6.2 | -6.1 | -6.1 | -6.62 |
| NavPaS | 11-deoxy-lactone-TTX | -8.3 | -6.8 | -6.8 | -6.7 | -6.6 | -6.5 | -6.4 | -6.2 | -6 | -6.70 |
| NavPaS | 1-hydroxy-8-epi-5,11-dideoxy-TTX | -7.2 | -6.6 | -6.6 | -6.2 | -6.1 | -6 | -6 | -6 | -5.9 | -6.29 |
| NavPaS | 4,9-anhydro-6-epi-TTX | -8 | -7.5 | -7.5 | -6.9 | -6.7 | -6.6 | -6.4 | -6.3 | -6.2 | -6.90 |
| NavPaS | 11-deoxy-TTX | -8.5 | -6.9 | -6.9 | -6.7 | -6.7 | -6.5 | -6.5 | -6.3 | -6.3 | -6.81 |
| NavPaS | 4-epi-TTX | -7.5 | -6.5 | -6.5 | -6.4 | -6.4 | -6.3 | -6.3 | -6.3 | -6.2 | -6.49 |

**SUPPORTING FIGURES**

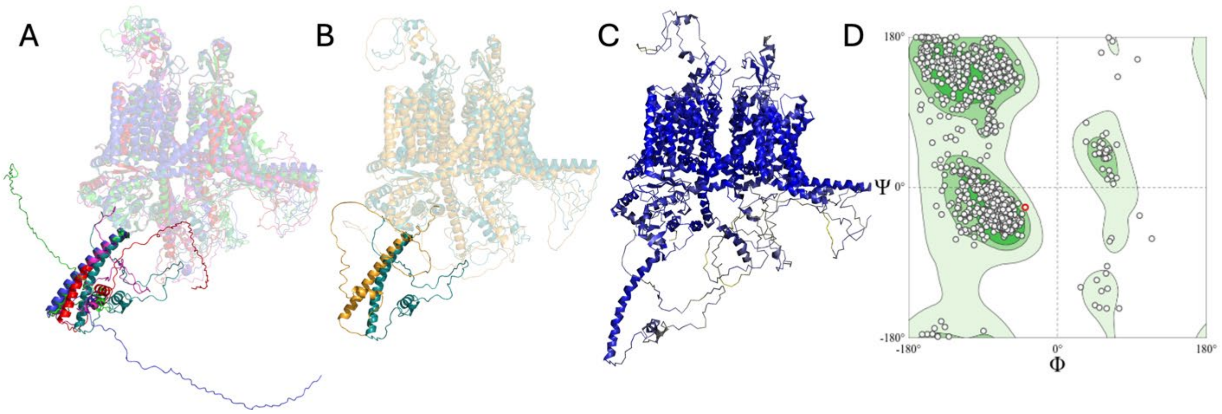


**Figure S1.** (A) All 5 models generated by Robetta aligned, highlighting the CTD to showcase 3 unusable models that have too much disorder for proper membrane insertion. The selected model is in teal. (B) Aligned AlphaFold Pufferfish NaV1.4a that was used as a template (yellow) aligned with the subsequently selected Robetta model (teal). (C) QMEANbrane scores displayed on the selected model (0.0 yellow to 1.0 blue). (D) Ramachandran for selected Robetta model.

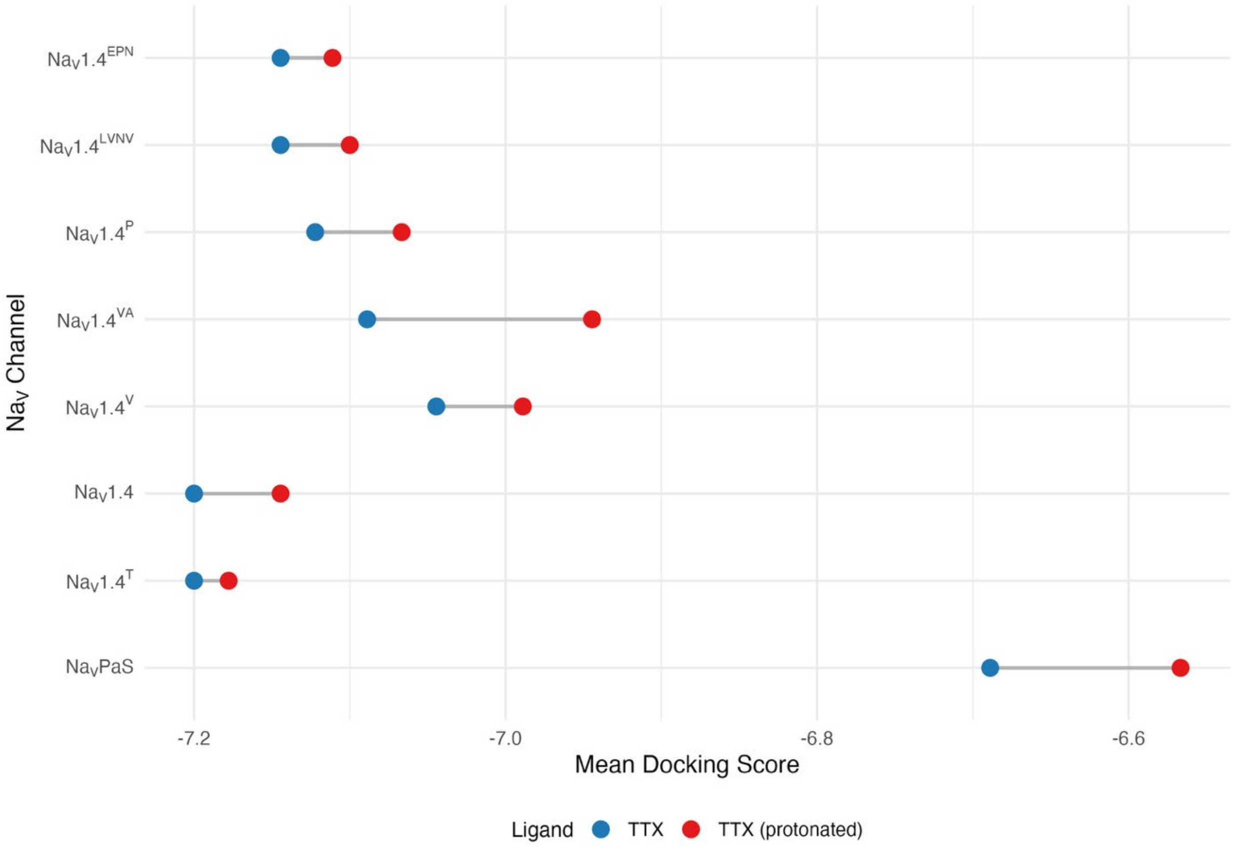


**Figure S2** Average docking scores for each of the seven snake $Na_V1.4$ channels and $Na_VPaS$ with canonical TTX (blue) and protonated TTX (red).

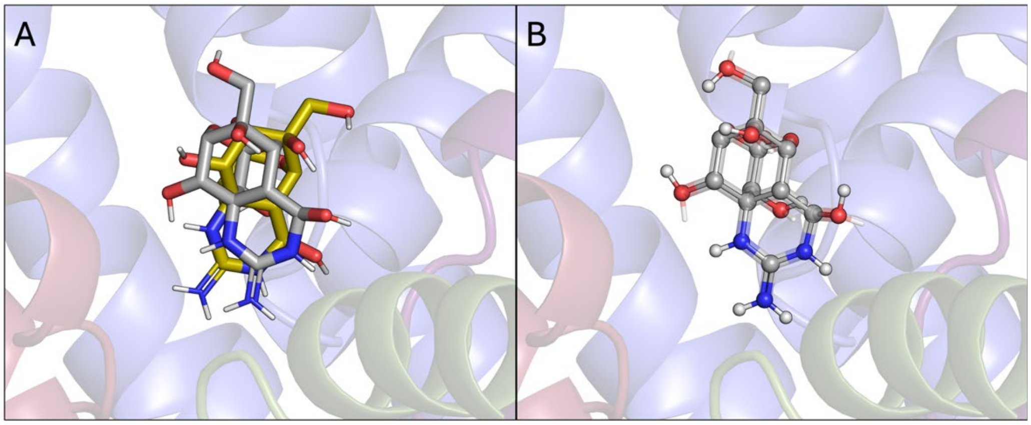


**Figure S3.** First poses of TTX in (A) $Na_V$PaS (gold) aligned with $Na_V$1.4 and (B) $Na_V$1.4 (transparent sticks) aligned with $Na_V1.4^{LVNV}$ (ball and stick).

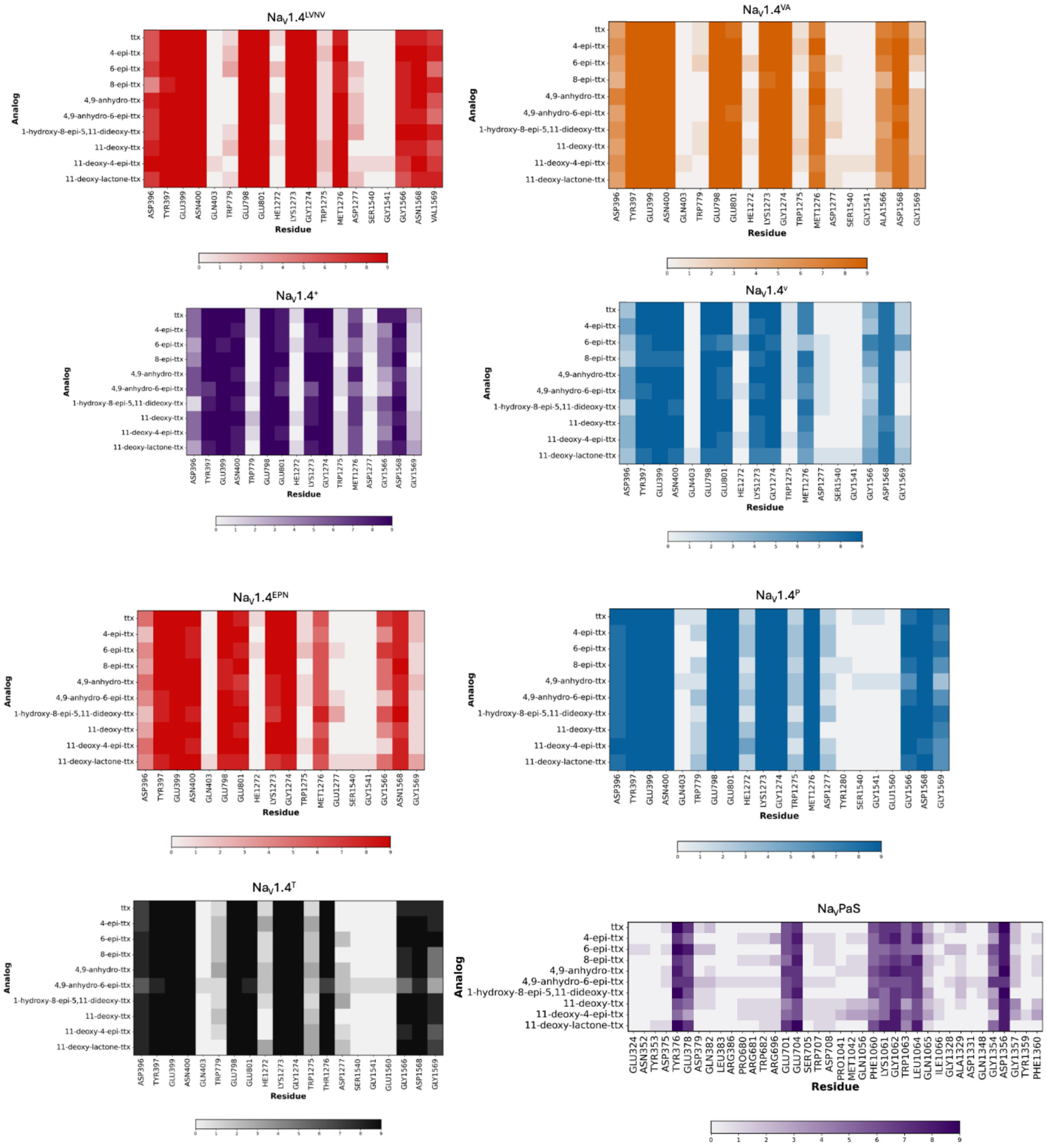


**Figure S4.** Interaction fingerprints across all analogs for all seven snake $Na_V1.4$ channel variants and the roach model $Na_VPaS$. Darker colors represent more poses (9 max) that have an interaction with that residue. Interactions include hydrophobic, aromatic, hydrogen bond donor, hydrogen bond acceptor, polar, charged, sidechain, and backbone.

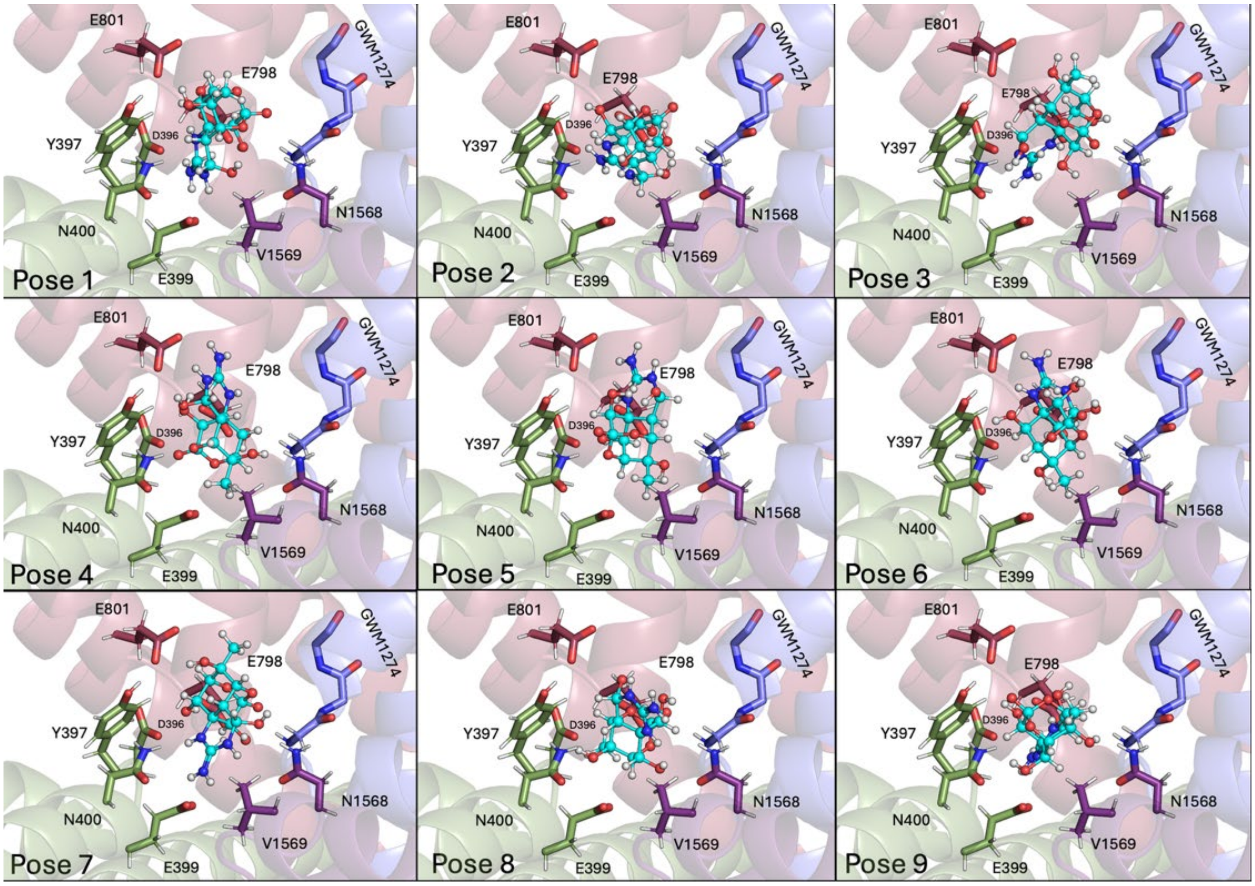


**Figure S5.** All nine poses for 11-deoxy-4-epi-TTX in $Na_V1.4^{LVNV}$ (cyan, colored by element). Some TTX-interacting residues are shown as sticks (colored by element) for reference.

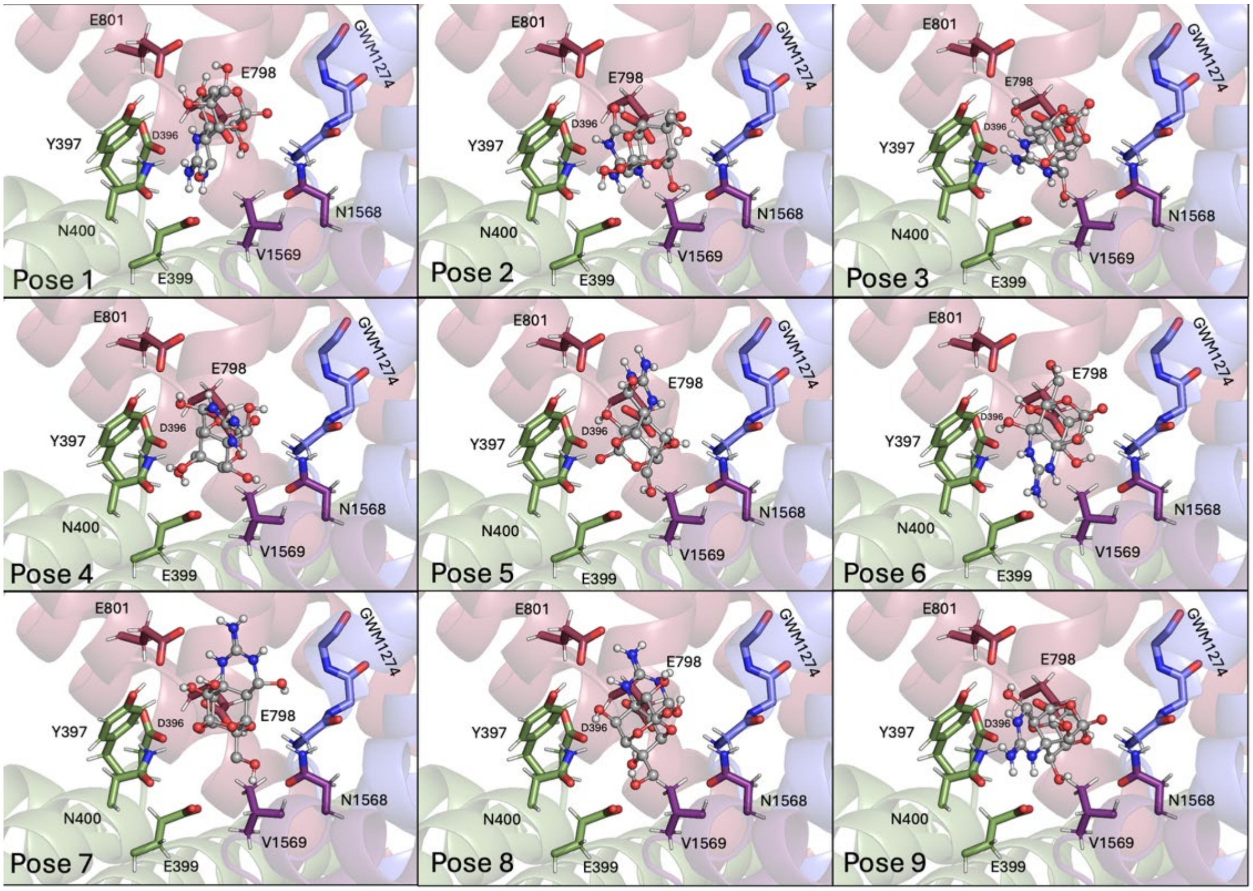


**Figure S6.** All nine poses for TTX in $Na_V1.4^{LVNV}$ (gray, colored by element). Some TTX-interacting residues are shown as sticks (colored by element) for reference.

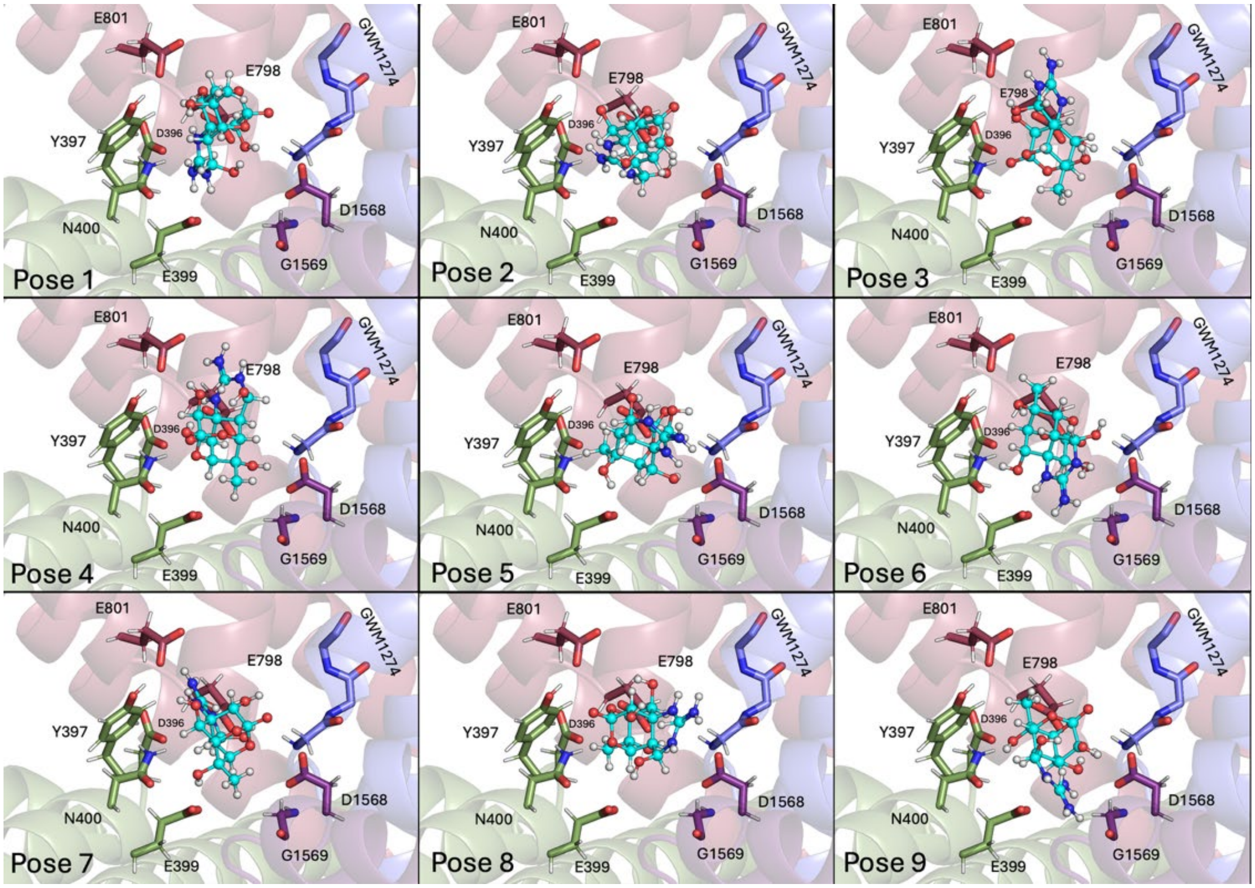


**Figure S7.** All nine poses for 11-deoxy-4-epi-TTX in $Na_V1.4$ (cyan, colored by element). Some TTX-interacting residues are shown as sticks (colored by element) for reference.

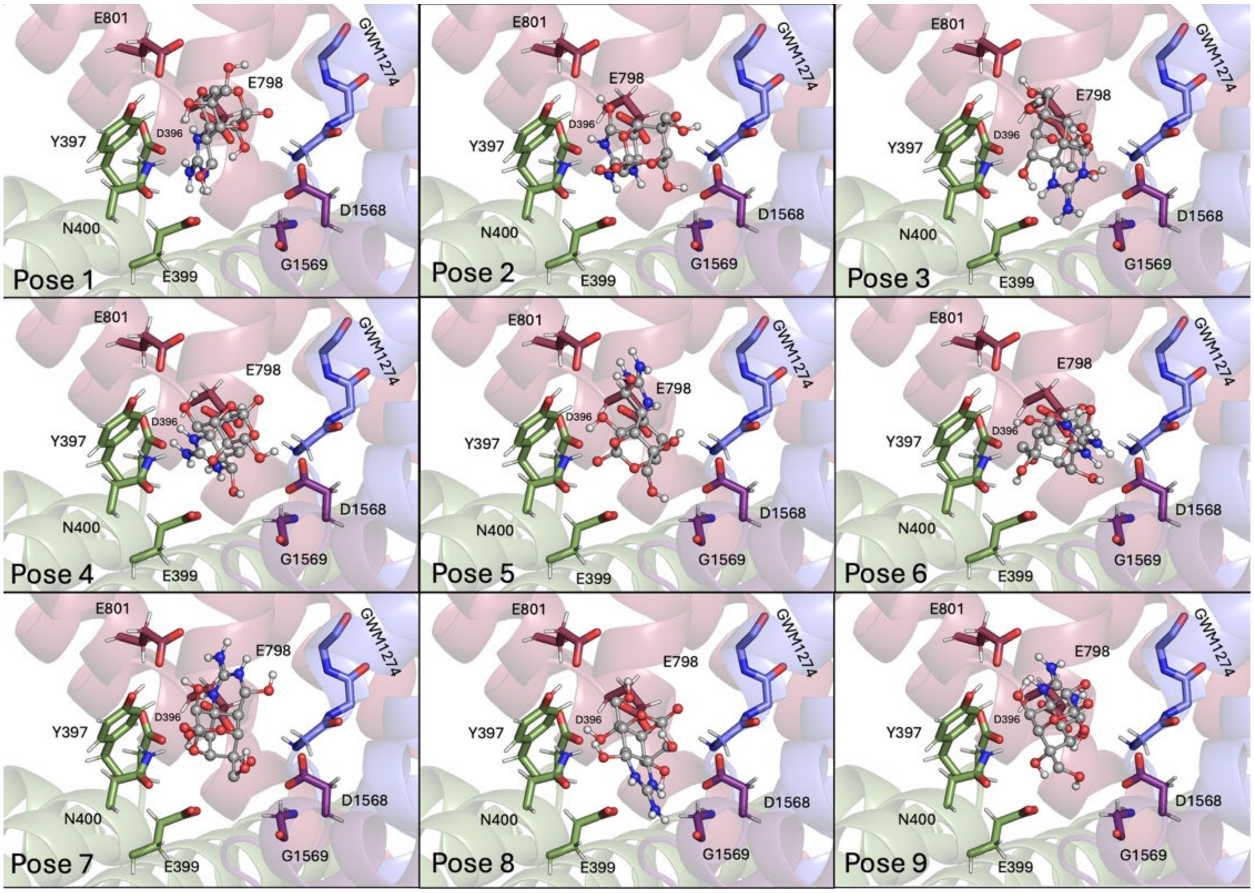


**Figure S8.** All nine poses for TTX in $Na_V1.4$ (cyan, colored by element). Some TTX-interacting residues are shown as sticks (colored by element) for reference.
